# Genetic and Cortical Cell-Type Liability Architecture of Autism

**DOI:** 10.64898/2025.12.10.693473

**Authors:** Thomas Renne, Florian Benitière, Cécile Poulain, Alma Dubuc, Vincent-Raphaël Bourque, Guillaume Huguet, Tomasz Nowakowski, Sébastien Jacquemont

**Affiliations:** CHU Sainte-Justine Azrieli Research Centre, Montréal, QC, Canada; Department of Biochemistry and Molecular Medicine, Université de Montréal, Montréal, QC, Canada; Department of Neurological Surgery, University of California, San Francisco, CA, USA; Weill Institute for Neurosciences, University of California, San Francisco, CA, USA; École Normale Supérieure de Lyon, Université Claude Bernard Lyon 1, Lyon, France; Program in Computational Biology and Bioinformatics, Yale University, New Haven, CT, USA; Division of Child Psychiatry, Faculty of Medicine, McGill University, Montréal, QC, Canada; Department of Pediatrics, University of Montreal, Montreal, Quebec, Canada

## Abstract

Autism Spectrum Disorders (ASD) can result from rare genetic variants interfering with brain development. Whether their effects converge on specific cortical cell types remains unresolved. Previous studies have focused on a narrow set of high-confidence ASD (hcASD) genes, which were enriched in neuronal cell types during prenatal development. By contrast, studies of postnatal cerebral cortex have repeatedly associated ASD with transcriptional changes in both neurons and glia.

To comprehensively map ASD genetic liability across cortical cell types, we conducted a functional genetic burden analysis with 124,416 individuals, including ASD probands and unaffected family members. We examined six classes of rare gene-disrupting variants aggregated across a complete spectrum of transcriptomic cell types of the human prefrontal cortex throughout development.

We show that cellular liabilities in ASD delineate a broad and developmentally dynamic architecture. Likewise, we uncover high dependency on classes of variants with Loss-of-Function (LoF) and *de novo* linked to prenatal cells, while duplications, missense, and inherited variants increase liability through postnatal and glial cell types. Notably, inherited LoF variants uncover the contribution of microglia to ASD liability, also supported by transcriptomic evidence from postmortem ASD brains. Finally, we show that overall, variants disrupting genes differentially expressed in postmortem ASD brains significantly contribute to ASD liability, demonstrating convergence between disrupted transcriptomes and genetic liability. Together, our study offers an integrative, cell-type-aware framework for interpreting ASD risk genetics.

## Introduction

Rare variants are major contributors to Autism Spectrum Disorders (ASD)^1^, and exome/genome sequencing studies have identified as many as 283 high-confidence ASD (hcASD) genes^2–4^. However, the vast majority of these genes were discovered by measuring an excess of *de novo* variants. This methodological approach could overlook ASD-associated genes that are not under extreme evolutionary constraint. Complementary studies have shown that rare inherited gene-disrupting variants also contribute to liability for ASD ^1,5–7^. It has been estimated that rare inherited and *de novo* LoF variants both account for the same population attributable risk for ASD^1,8^, and that 80% of ASD risk conferred by rare inherited variants do not disrupt known ASD-associated genes^1^. This diversity of variants and genes involved raises the question of whether they converge on shared cellular pathways.

Indeed, cell types are the fundamental unit of brain architecture. Therefore, understanding how ASD-associated genes and genetic variants might disrupt their function represents a central goal in neurodevelopmental research^9–11^. Prior studies highlighted enriched co-expression of hcASD genes among glutamatergic excitatory neurons and GABAergic interneurons^2,3,5,12–14^. However, because most of those efforts utilized transcriptomic data from the prenatal stage, they may not fully capture the associations of postnatal cell types with ASD.

Unexpectedly, these predicted cellular liabilities remain discordant with transcriptomic dysregulation in cell types observed between ASD *and* non-ASD postmortem brain tissue. In fact, the highest number of differentially expressed (DE) genes^15,16^ have been observed in astrocytes, microglia, oligodendrocytes, and deep-layer excitatory neurons. This discordance between genetic associations and postmortem findings could be partly explained by the different developmental epochs tested with these two approaches, hence arguing the need for a cross-developmental analysis.

Finally, current gene association methods rely on gene-level burden tests, which aggregate all classes of gene-disrupting variants. This gene-centric approach may obscure potential differences in how distinct variant classes contribute to ASD risk and affect specific cell types^17^. Previous work has shown that deletions and duplications (constituting opposite gene dosage deviations^18^) alter distinct biological processes^19–22^, and emerging proteomics studies have begun to decipher the heterogeneous effects of protein-altering variants^23^.

In summary, critical gaps persist in our understanding of cellular liabilities linked to the broad spectrum of gene-disrupting variants contributing to ASD. To address this question, we developed a Functional Burden Association Test (*FunBAT,* adapted from previously published burden approaches^19,24,25^) which aggregates, across 18 cortical cell types over 6 developmental epochs, gene-disrupting variants in 124,416 individuals, with ASD and their unaffected family members. Beyond the known prenatal cell-type associations with ASD, we identified cellular liabilities for ASD at postnatal epochs and within glial cell types. These cell-type liabilities were specific to classes of genetic variants and mapped to broad ASD subgroups (i.e., early and late diagnosis). Our results bridge genetic association and postmortem transcriptomic studies, specifically that ASD-DE genes also contribute to ASD risk when disrupted by rare variants. Finally, while functional burden does not provide gene-level association with ASD, gene prioritization highlighted glial marker genes as potential contributors to ASD liability.

## Results

### Quantifying cell-type specificity across coding genes

Previous studies investigating cell-type enrichments of hcASD genes have employed a broad range of methods without clearly quantifying the tradeoffs between cell-type sensitivity and specificity. To identify cell-type-specific genes, we first quantified the expression specificity and sensitivity of 19,152 autosomal protein-coding genes across cell types and developmental stages of the cerebral cortex. We leveraged single-nucleus RNA sequencing (snRNAseq) data ^26^ on 153,325 cells (26 prefrontal cortical samples from gestational week 22 to 40 years old), defining 18 cell types across 6 developmental periods: prenatal, neonatal, infancy, childhood, adolescence, and adult (**Figure 1.A**).

**Figure 1.**
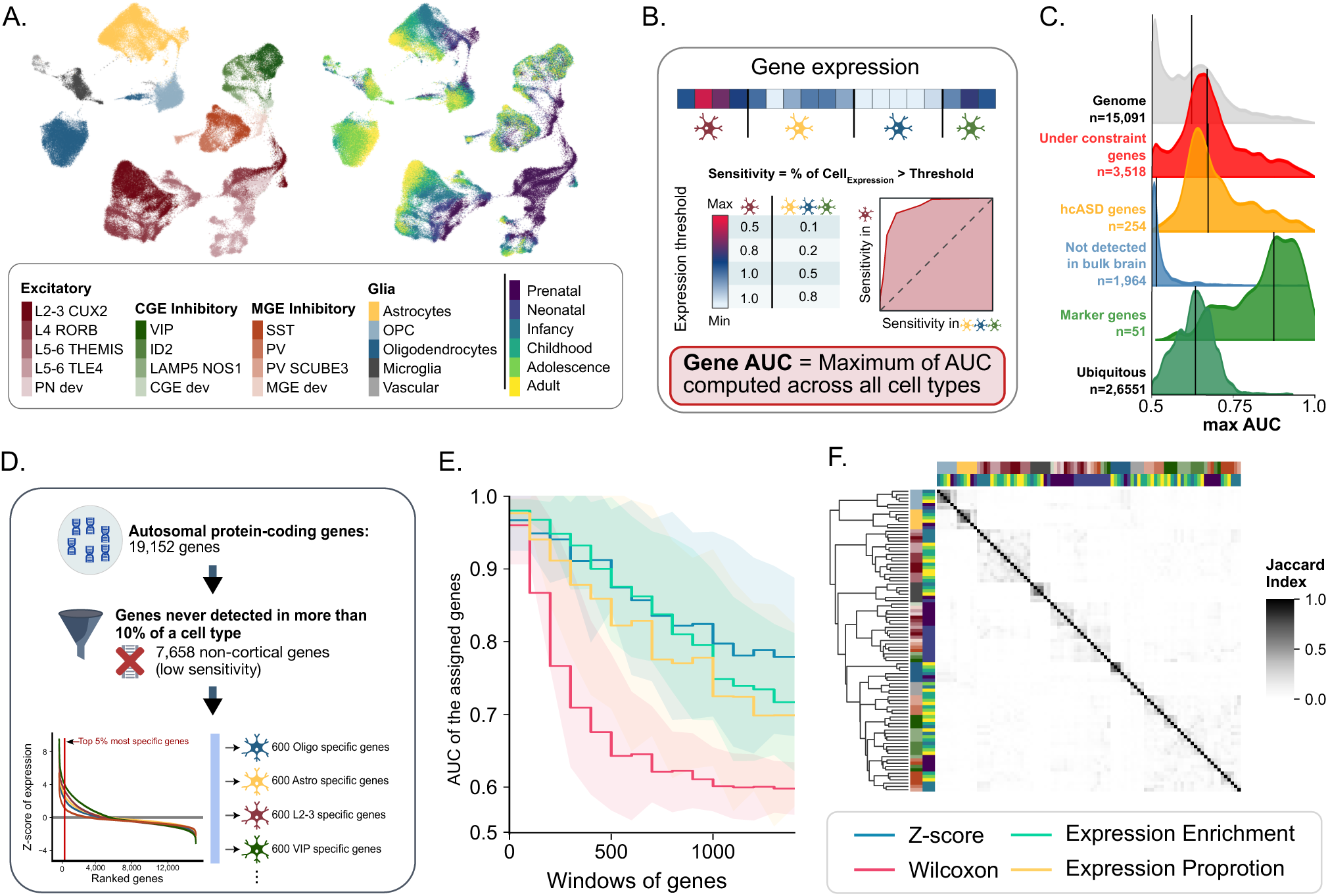
Defining and characterizing cell-type-specific gene sets. (A) UMAP of the 153,325 cortical cells divided into 18 manually annotated cell types and 6 developmental epochs. (B) Area Under the Curve (AUC) quantifying a gene’s sensitivity and specificity in discriminating a given cell type. (C) Distribution of maximum AUCs for subsets of genes characterizing their ability to discriminate between 91 cell types and 6 developmental stages. From top to bottom distribution: all autosomal protein-coding genes, genes under genetic constraint (LOEUF_V4 <= 0.6)^34^, hcASD genes, genes not expressed in the brain^55^, marker genes used to annotate the cell types, and genes with a ubiquitous expression^56^. In the figure, the number of genes refers to those identified in the snRNAseq dataset. (D) Method used to assign the most specific genes to each cell type based on their expression profile. (E) Performance (in discriminating cell types) of 4 methods used to create cell-type-specific gene sets. Y-axis: AUC of each gene set. X-axis: windows of 100 genes from the highest to the lowest specificity. (F) Gene overlap across all cell-type-specific gene sets. The color code represents the Jaccard index.

We computed the fraction of cells expressing a gene within (sensitivity) and outside (1 - specificity) each cell type cluster (**Supp figure 1, A,B**). The trade-off between both metrics is represented by the Area Under the Receiver Operating Characteristics Curve (AUC) and indicates how effectively the expression of a specific gene can distinguish one cell type from others. (**Figure 1.B**). The 52 marker genes used to annotate cell types had a high median AUC_max_ = 0.87 (AUC_max_ i.e., the highest discrimination value across all cell types; **Figure 1.C**). In contrast, previously published hcASD genes (n = 283, **Supp figure 2.A**) showed lower specificity and AUC_max_ when compared to all genes expressed in the cerebral cortex. (medians = 0.60 and 0.87, p-value = 7e^-51^; **Figure 1.C, Supp Figure 1.B-C**), consistent with the fact that many ASD risk genes are broadly expressed across brain cell types[27]. Lower specificity was also observed for intolerant genes (**Supp Figure 1.D**).

### Functionally defined gene sets accurately predict cell-type clusters

We explored the performance of 4 methods^28^ used to define cell-type-specific gene sets: z-score^19^, Wilcoxon test, Expression Proportion (EP)^29,30^, and Expression Enrichment (EE)^31^, (**Figure 1.D**). We characterized the ability of gene sets derived from these methods to discriminate one cell type from another (**Figure 1.E**). Gene sets defined by the z-score approach showed the highest AUC (Student’s t-test FDR = 1e-4 to 1e-274) (i.e., the best tradeoff between cell-type-specificity and sensitivity) while limiting the number of genes never assigned to any cell type (c.f., method). We therefore employed this method for all subsequent analyses, using a gene-set size of 600 genes, achieving an AUC > 0.9. Overall, cell-type-specific gene sets showed limited overlap with a mean Jaccard index of 3%. (**Figure 1.F, Supp figure 3.A-C**). Of note, the created gene sets were more similar (higher overlap) within each cell type than within each developmental stage. Nonetheless, our developmental gene sets were in line with previously published prenatal and postnatal trajectories of gene expression measured during brain development^32^ (**Supp figure 4**).

### hcASD gene enrichment among cell types and developmental epochs

Consistent with prior studies ^2,3,5,13^, hcASD genes were enriched in both excitatory (L2-3 CUX2, L4 RORB, and L5-6 TLE4) and inhibitory prenatal neurons (VIP, ID2, SST, PV_SCUBE3 and MGE_dev, **Figure 2.A**). We also observed enrichment for three prenatal glial cell types (astrocytes, oligodendrocytes, and OPC) and many postnatal (infancy) cell types, which was not previously established^33^. Sensitivity analyses showed that modulating the level of cell type specificity of gene sets (which also changes gene set sizes) did not influence these findings (**Figure 2.B, C**).

**Figure 2.**
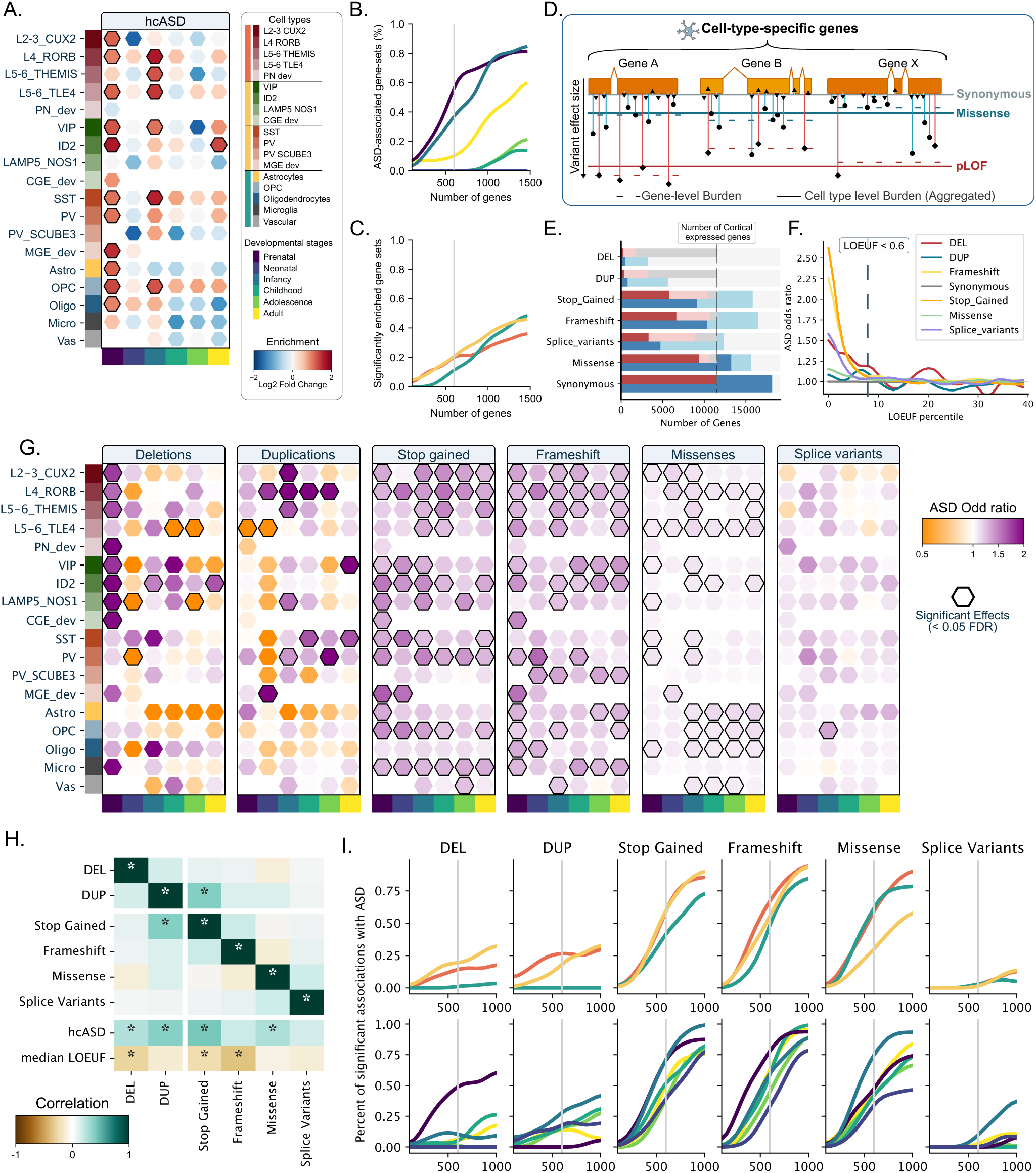
Functional genetic burden associates a broad range of cell types with ASD. (A) Enrichment of 283 hcASD genes across 18 cell types and 6 developmental epochs. Positive and negative enrichments are represented in red and blue, respectively, and significant enrichments are outlined in black. Rows represent 18 different cell types, and columns represent 6 developmental epochs. Percentage of significant positive enrichment of the 283 hcASD genes among the 3 main cell classes (excitatory neurons, inhibitory neurons, and glia) (B) and the 6 developmental epochs (C) for 15 levels of specificity (i.e., from the top 100 most specific to the top 1,500 most specific in each cell type). Curves smoothed with a Gaussian filter with a sigma = 2. (D) To overcome the power issue of the gene-level association tests for rare variants, FunBAT aggregates all the gene-disrupting variants assigned to the same cell type and computes its average liability (functional burden) for each variant class. (E) Number of autosomal genes (red: cortically expressed and blue: expressed in any tissue) disrupted by at least one (light) or ten (dark) rare variants across 141,136 SPARK individuals. (F) ASD liabilities computed for 40 windows of genetic constraint when disrupted by each of the 7 types of variants. Significant associations (FDR < 0.05) are represented with larger points. **(G)** ASD liability computed for 91 cell-type gene sets across 6 types of variants. Represented liabilities depicted the under-genetic-constraint subset of genes (LOEUF < 0.6). Increased and decreased risk for ASD are represented by purple and orange hexagons. Significant associations are outlined in black (FDR < 0.05). (H) Matrix representing pairwise correlations between effect sizes for the 91 cell-type liabilities across 6 classes of variants, as well as enrichment of high-confidence ASD genes (hcASD) and the median level of constraint computed for all 91 cell types (LOEUF). The color scale represents the Pearson r-value, and stars depict significant correlations. **(I)** The number of cell types associated with ASD increases as gene sets are less specific and larger. The Y-axis represents the percent of significant association with ASD for the main cell classes (top) and the developmental stages (bottom). The X-axis represents the size of the gene sets ranging from the top 99th (n=100) to the top 90th (n=1100) percentile of genes with the highest relative expression in a given cell type.

### A broad landscape of cell types is implicated in ASD

To investigate cell-type liabilities beyond known hcASD genes, we developed a Functional Burden Association Test (FunBAT), which aggregates all rare variants disrupting genes assigned to a given cell type to compute a burden association with ASD (**Figure 2.D**). We implemented burden tests in 124,416 SPARK individuals (**Supp Table 1**) across 7 classes of variants (deletions, duplications, stop-gained, frameshifts, missenses, splice variants, and synonymous variants covering 15% to 100% of the cortically expressed genes, **Figure 2.E**) and across 40 levels of genetic constraint^34^. In line with previous studies^1^, we showed that all genetic associations with ASD were observed for genes under genetic constraint (**Figure 2.F**). We thus implemented *FunBAT* across 91 cell-type gene sets, restricting our analysis to the 3,784 genes with LOEUF_V4 below 0.6 (total of 637 association tests). Sensitivity analyses did not identify any significant associations with ASD for subsets of cell-type-specific genes under less genetic constraint (LOEUF >= 0.6) or for non-brain-expressed genes **(Supp figure 5 and 6**).

### Cell type liability for ASD is dependent on classes of genetic variants

Significant associations with ASD were observed in 0% of cell type gene sets for splice variants and negative controls (synonymous) to over 50% of cell types for stop-gained and frameshifts (**Figure 2.G**). Each class of gene-disrupting variant showed distinct patterns (maps) of cell type liability (**Figure 2.G,I**) with mild to no correlation between effect-size maps computed for the 6 classes of gene-disrupting variants (median rs = 0.06). The correlation was also low between stop-gains and frameshifts (Spearman rs = 0.17). This was not explained by the frequencies of these two classes of loss-of-function variants, which were similar across the different gene sets and genes (rs = 0.94 and 0.76, **Supp Figure 7**). Some cell-type liabilities were specific to classes of variants, such as PV SCUB3 and SST inhibitory neurons, which showed ASD liability only when their assigned genes were disrupted by frameshift and stop-gain, respectively, while microglia were only associated with ASD for LoF variants.

Prenatal cell types were a common denominator across loss-of-function (LoF) variants (deletions, stop-gains, frameshifts), with ASD-associated deletions showing preferential liability for prenatal excitatory and CGE-derivated inhibitory neurons, while stop-gains and frameshifts showed associations across most neuronal and glial prenatal cell types. Contrary to LoF variants, duplications and missense variants primarily implicated postnatal cell types. ASD-associated missenses were linked to postnatal excitatory neurons, astrocytes, and OPC, while duplications showed associations through postnatal excitatory (in particular, cortical layer 4) and inhibitory neurons. Sensitivity analyses showed that increasing or decreasing the level of cell-type specificity of the gene sets (therefore changing gene set size) did not influence these results (e.g., prenatal cell types and glial cells remained the strongest association for LoF and missense variants, respectively, regardless of gene-set specificity and size; **Figure 2.I**). We showed that these cell type liability patterns required a sample size of approximately 100,000 to become stable and robust (i.e., Spearman rs >= 0.8). Specifically, we recomputed the burden association tests in smaller and smaller randomly sampled subsets of the genetic dataset (**Supp Figure 8**). As an example, the novel microglial association with ASD was detected with a power of 80% for sample sizes above 110,000.

### Cell types linked to *de novo*, inherited, and novel ASD genetic contribution

To understand cell-type liabilities linked to novel ASD genetic contributions, we excluded 283 hcASD genes from our analyses. The resulting cell-type liability maps were highly correlated (rs = 0.81 to 0.99, **Figure 3.A**) with the initial maps with a small drop in effect sizes (0 to 7%), suggesting that many cell-type liabilities for ASD described above were not solely related to hcASD genes. Notably, microglia, LAMP5, CGE precursor inhibitory neurons, and more generally postnatal cell types are likely linked to novel genetic contributions to ASD (**Figure 3.B**). We asked if these novel cell-type liabilities were related to *de novo versus* inherited variants. To this end, we identified *de novo* variants in 31,786 trios (22,322 ASD and 9,464 control siblings). The strongest enrichments of *de novo* variants (ASD versus control) were observed in excitatory neurons and the prenatal stage (**Supp Figure 9**). In contrast, among autistic individuals, *de novo* variants were depleted in glial-specific gene sets (relative to the rest of the brain-expressed genes) as well as postnatal cell types with a gradual decrease throughout development (**figure 3.C, Supp figure 10**). Finally, we recomputed cellular liabilities for ASD after removing de novo variants. Although the significance of the associations with ASD diminished considerably, the estimates across cellular liabilities remained strongly correlated (rs = 0.73 to 0.93, **Figure 3.D**), with liability estimates for excitatory neurons and glia being the most and the least impacted by this removal, respectively.

**Figure 3.**
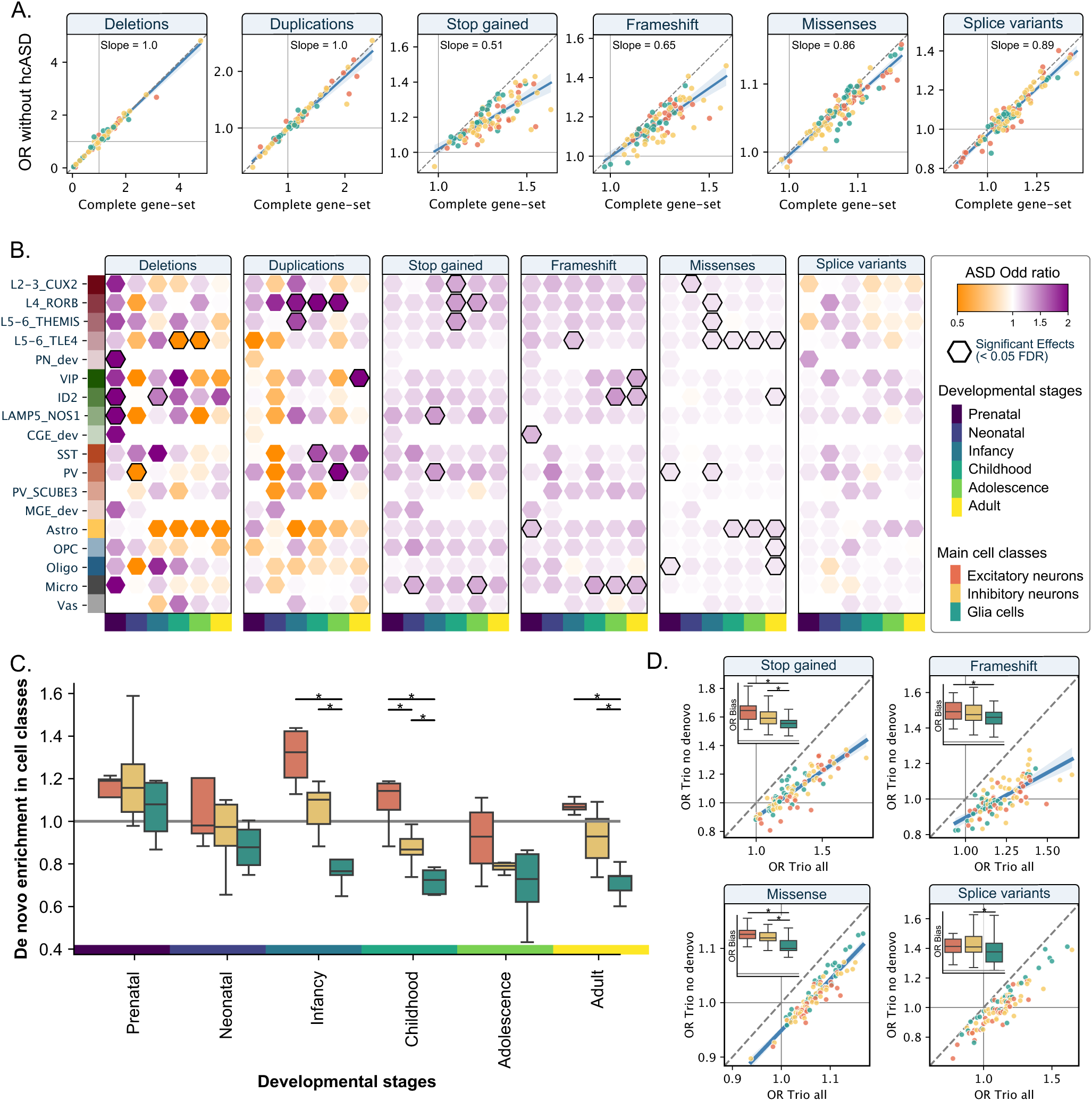
Cell-type liability for ASD beyond the hcASD genes. (A) Concordance analyses of ASD liabilities before (X-axis) and after (Y-axis) excluding 283 hcASD genes. Each point is the estimate (Odd Ratio) for ASD liability for each of the 91 a cell types. Color-codes represent the 3 main cell classes. The diagonal dashed line represents the exact concordance. (B) ASD liability computed for 91 cell-type gene sets, after excluding 283 hcASD genes. Color scale represents the Odds Ratio. Significant associations are outlined in black (FDR < 0.05). (C) Enrichments of *de novo* short variants in ASD individuals across 91 developmental cell types (**Supp Figure 9**) grouped in 3 main cell classes. Stars between classes indicate significant differences (FDR < 0.05) in the enrichment levels. Values above and below 1 represent cell types enriched and depleted in de novo variants relative to all other brain-expressed genes, respectively. (D) Concordance analyses of ASD liabilities before (X-axis) and after (Y-axis), excluding de novo variants in 31,814 trios. Boxplots represent the biases (residuals) observed for 3 classes of cell types after removal of de novo variants. Lower values indicate liabilities less impacted by the de novo removal. Significant differences between the biases computed for the 3 main classes are represented by stars.

### Genes contributing to cell type liability for ASD

We asked whether the cell types associated with ASD were due to disrupting variants broadly distributed across a given gene set or, instead, to a smaller number of variants (sparse alternative). To do so, we compared results from the SKAT-Burden and SKAT-Variance methods^35^, which are sensitive to the broad and sparse hypotheses, respectively. Overall, burden tests provided many associations between rare variants and ASD, while SKAT-Variance did not detect any (**Figure 4.A**). Cell type liabilities with the strongest bias towards the sparse alternative were those the most enriched in hcASD genes (e.g., microglia, rs = 0.51 to 0.74; p-value = 2e-7 to 3e-17).

**Figure 4.**
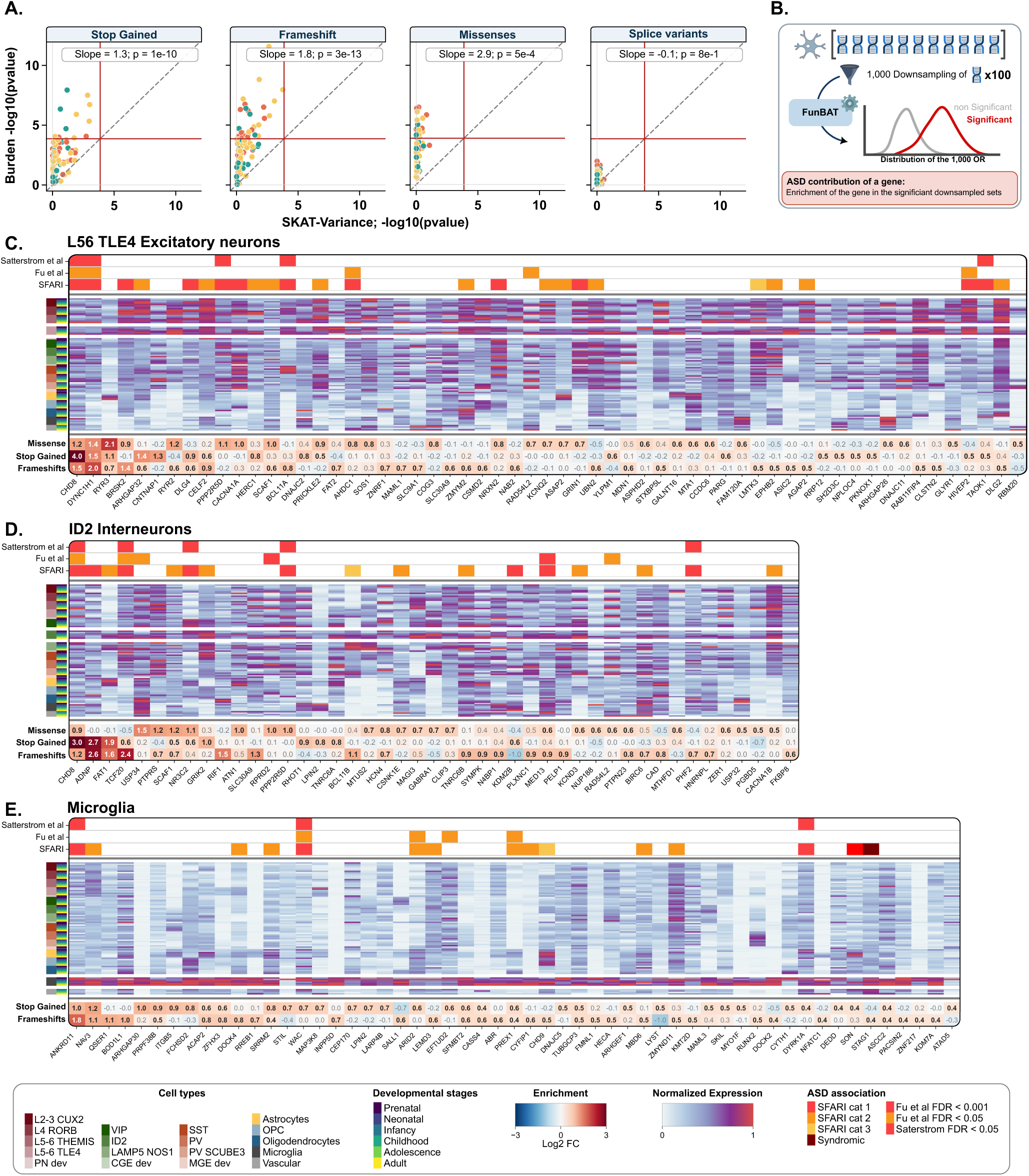
A broad set of genes contributes to cellular liabilities for ASD. (A) Concordance analysis between the p-values (-log10) of the associations provided by SKAT-Variance^35^ (sparse hypothesis) and SKAT-Burden (broad hypothesis). Data points correspond to the 91 cell types. The bias is reported for each class of variants as the slope of the regression line. (i.e., a slope of 3 indicates that p-values are 1,000 times smaller). The red lines represent the thresholds of significance (p=1e-4; Bonferroni correction, n = 364 tests). (B) Down-sampling method used to prioritize genes contributing to significant burden associations between cell-type gene sets and ASD. All the genes of the cell type of interest are randomly down-sampled (100 genes) one thousand times. Top (significantly enriched in significant down-sampled sets) contributing genes are prioritized (from left to right) for (C) L56_TLE4 excitatory neurons, (D) ID2 inhibitory neurons, and (E) Microglia. Previous ASD associations of the genes are color-coded on the top for 3 different resources: SFARI^4^, Satterstrom et al.^3^ and Fu et al.^2^ In the middle is represented the normalized expression (MinMax scaling for each gene) across the 91 cell types for each gene. The enrichment levels (log2) for the prioritized genes are shown for the different classes of variants. Significant enrichments are in bold and black.

We sought to rank/prioritize the contribution of genes for 3 cell types (one from each cell class) that remained associated with ASD after removing known hcASD genes (i.e., L5/6-TLE4 excitatory neurons, ID2 inhibitory neurons, and microglia). To do so, we conducted burden association tests on downsampled gene sets by adapting a previously published method^36^ (**Figure 4.B**). For the three cell types, 13%, 12%, and 17% of their genes (under constraint) were significantly enriched in ASD-associated downsampled gene sets, respectively, and therefore prioritized as candidate genes driving the association with ASD (**Figure 4.C-E**). The vast majority of ASD-candidate genes for the microglia (>90%), excitatory and inhibitory neurons (80%) were not previously reported as hcASD genes. For microglia, many candidate genes were highly cell-type specific (including well-established cell-type marker genes such as *SALL1, DOCK4, BOD1L1* **Figure 4.E**). In contrast, for ID2 GABAergic interneurons and deep-layer excitatory neurons, the ASD candidate genes were more broadly expressed across several cell types (**Figure 4.C,D**). This was in line with previous studies identifying these two cell types as enriched in hcASD genes^2,3,5^. Finally, the 55 candidate genes identified in microglia were significantly enriched in the nucleoplasm and in the RHO GTPase cycle, while gene enrichments of the two neuronal cell types were associated with synapse and somatodendritic compartment and molecular functions, in line with previous findings^11^ (**Supp Table 2-4**).

### Genes differentially expressed in ASD brain cell types contribute to ASD liability

Previous studies of transcriptomic changes in ASD brains have repeatedly identified increased abundance of activated microglia and astrocytes, with patterns of DE genes reported across most cell types^15,16,37^. However, these cellular ASD-DE genes show little overlap (4 to 12%) with known hcASD genes, which (**Figure 5.A**). We first sought to understand the general characteristics of ASD-DE genes. As expected, they were enriched in postnatally (p-value = 1e-11) and depleted in prenatally expressed genes^32^ (p-value = 1e-2). Both down- and up-regulated genes exhibited higher intolerance to haplosufficiency compared to the whole protein-coding genes as well as the cortically expressed genes examined above (p-value = 1e-103 and 2e-75**, Supp figure 11**). ASD-DE genes also showed low cell-type specificity and limited overlap (12% and 20% for up- and down-regulated genes) with their corresponding cell-type-specific genes (i.e., ASD-DE genes in microglia were not microglial-specific genes, **Figure 5.B**). Such discordance may be explained by our findings showing that ASD-DE genes (both up- and down-regulated) were only enriched in postnatal rising genes (OR = 0.4 and 0.4, p-values = 1e-11 and 9e-11 *versus* prenatal falling genes ORs = -0.18 and -0.05, p-value = 1e-2 and 4e-1). The level of enrichment in their corresponding cell types also increased along the developmental stages (**Supp Figure 12**), whereas hcASD genes were more prenatal, as demonstrated above. We then used *FunBAT* to test the association between ASD and the burden of variants disrupting 47 sets of genes previously identified as up- or down-regulated in brain cell types of ASD individuals^15^. The majority (76%) of these ASD-DE gene sets were significantly associated with ASD when disrupted by at least one class of rare variants. Even after removing hcASD genes (4% of the whole ASD-DE genes), 34% of gene sets remained significantly associated with ASD, demonstrating that ASD-DE genes in postmortem ASD brains also contribute to liability for ASD when disrupted by rare variants. Down-regulated gene sets showed a larger number of significant associations compared to up-regulated gene sets across all classes of variants, and this, in an uncorrelated way (29% versus 9%; **Figure 5.A,C,D**) and higher OR after correcting for the gene-set size (**Figure 5.E**). As shown above, there was limited overlap between cell-type-specific and the ASD-DE genes. As a result, we observed no correlation between ASD liabilities computed using cell-type-specific genes versus ASD-DE genes (**Figure 5.D**). Despite this lack of concordance, 3 cell types were associated with ASD when burden association was computed with either cell-type-specific genes, up- or down-regulated genes (i.e., OPCS_1, EXT_6_L23, EXT_4_L56, **Figure 5.C**).

**Figure 5.**
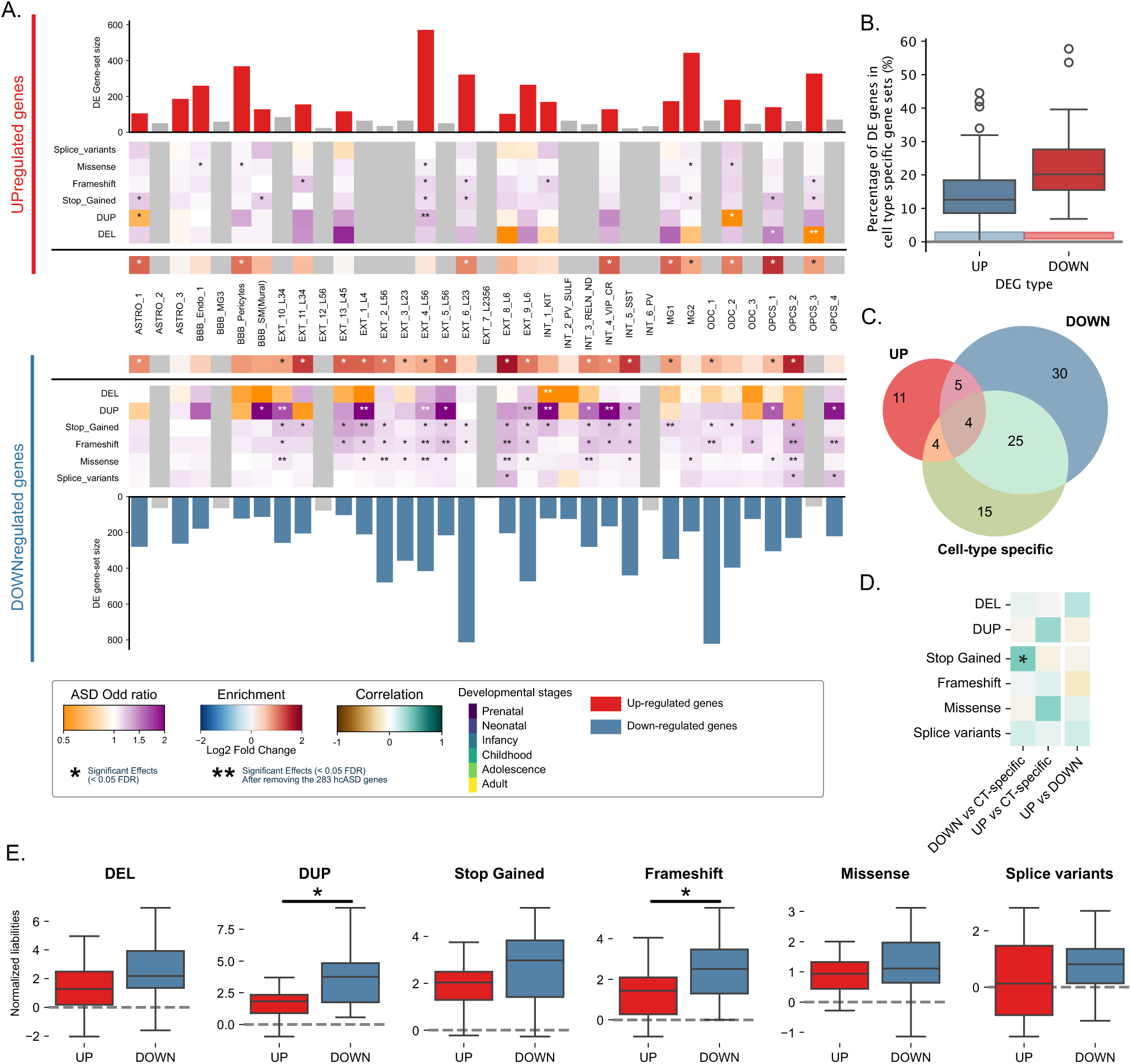
Effects of variants disrupting genes differentially expressed in the cerebral cortex of ASD individuals. (A) ASD liabilities computed for sets of differentially expressed (DE) genes across 30 cell types. The top and bottom bar plots show the number of up- and down-regulated genes identified in ASD prefrontal cortices from Wamsley et al.^15^ respectively. The gray bar plots are cell types with less than 100 DE genes, not included in our functional burden analyses due to limited statistical power. The % of hcASD in each cell-type DE gene set is shown at the center of the figure. The orange-to-purple matrices depict the association (odd ratio) with ASD for each DE gene sets when they’re disrupted by 6 types of variants. FDR significant burden associations (47 DE gene sets x 6 variant classes) are labeled with a star. ASD liabilities with double stars indicate that the burden test remained significant after removing the 283 hcASD genes. (B) Boxplots comparing ASD liabilities between up- and down-regulated gene sets (normalized for gene set size) Stars indicate significant differences in liabilities between the up- and down-regulated gene sets. (C) Venn diagram representing the overlap between the 216 (36 cell types x 6 variant types) cell-type liabilities for ASD when computed using cell-type specific gene sets, up- and down-regulated genes. The 4 overlapping association are missense of OPCS_1, frameshift and stop-gained of EXT_6_L23, and frameshifts of EXT_4_L56 (D) Correlations between patterns of ASD liabilities computed using cell-type specific gene sets, up- and down-regulated genes. Positive correlations are depicted in green; the significant correlations are labeled with stars. (E) Spearman correlation of the ASD liabilities computed for the up- and down-regulated gene sets against the ASD liabilities computed for the corresponding cell-type-specific gene sets.

### Cell type liability and ASD clinical heterogeneity

We asked if the relationship between inheritance, classes of variants, and cell type liabilities for ASD described above may relate to the clinical heterogeneity of ASD. We explored biological sex and age at diagnosis (before and after 4 years of age) ^38,39^, which are two major features associated with the clinical presentation and prevalence of ASD^40^ (1F:3.5M in our dataset). Consistent with prior observations^41,42^, we found more *de novo* stop-gained variants in females compared to ASD males (Mann-Whitney U test, p-value = 5e-3, **Supp Figure 13**) as well as more hcASD disrupted by LoF and missenses(Fisher exact test, OR = 1.29 and 1.27; p-value = 8e-6 and 5e-7). Contrary to our expectations^43^, we found no overall differences in the de novo rates between the early and late diagnosis groups, and such for the two sexes (**Supp Figure 14 and 15**). Despite these differences in *de novo* variants, we did not identify any broad male/female or early/late diagnosis biases for the level of ASD liability along the main cell classes or the developmental epochs (**Supp Figure 16**). In other words, variants disrupting genes of a given cell type were associated with the same level of ASD liability for both sexes and both age-at-diagnosis groups.

## Discussion

By combining a comprehensive single-nucleus transcriptomic atlas and a functional burden approach (FunBAT), we systematically mapped cell-type liabilities for ASD. We identified novel ASD-associated cell types, in particular, PV_SCUBE3 inhibitory neurons, microglia, OPCs, and postnatal cells, which were depleted in de novo (compared to inherited) variants, explaining why they had eluded detection in previous studies. The enhanced power provided by FunBAT enabled us to demonstrate that patterns of ASD-associated cell type liability were specific to classes of variants: loss-of-function preferentially mapped to prenatal cell types, while missenses and duplications were linked to postnatal excitatory neurons, astrocytes, and OPCs. Some cell types were uniquely linked to ASD through one class of variants, such as PV-SCUBE3 neuron liability, observed only for frameshifts. These cell type liability patterns were generalizable across males and females, as well as individuals with an early *versus* late ASD diagnosis. Finally, we demonstrate that ASD-DE genes represent distinct mechanisms and are mostly non-overlapping with cell-type-specific genes, but they significantly contribute to ASD when disrupted by rare variants.

We reveal distinct patterns of cell type liability to ASD across different classes of genetic variants and developmental epochs. Using the first uniformly sequenced RNA dataset of pre- and postnatal cortical nuclei^26^, we show that gene-disrupting variants were broadly distributed across cell types and developmental epochs and were mostly unrelated to hcASD genes. This is in line with previous studies suggesting that up to 70% of the inherited gene-disrupting variants contributing to ASD liability remain undocumented^7^. This broad spectrum of cell types and developmental epochs implicated in ASD contrasts with earlier genetic studies, which primarily reported prenatal neuronal vulnerability^2,3,32^. Several factors may explain this discrepancy. First, earlier genetic association studies of ASD often had lower statistical power due to smaller cohorts and gene-level burden methods. Second, they focused predominantly on *de novo* variants. We demonstrate that such variants under extreme selective constraint preferentially map onto prenatal developmental processes (in line with previous research^32^) and are depleted in postnatal. However, the lack of consensus on cell type definition and methods for assigning genes to cell types still hinders cross-study comparisons.

Prior genetic studies of rare variants have adopted a gene-centric approach, aggregating at the gene level all types of gene-disrupting variants with an emphasis on *de novo* occurrence^2,3,7,44^. However, our findings reveal distinct patterns of cell-type liability from one class of disrupting variant to another. This suggests that different variant classes implicate distinct biological processes in ASD liability, a pattern previously observed for deletions and duplications associated with neurodevelopmental conditions^19–22^. Notably, even complete gene deletions (gene dosage), stop gains, and frameshifts exhibited distinct patterns of cell-type liabilities, despite their presumed similarity in molecular consequences. This is in line with a growing number of observations demonstrating that genes show differential sensitivity to pure haploinsufficiency (gene dosage) *versus* stop-gained that may lead to truncated proteins with distinct deleterious effects ^45^.

The microglial alterations repeatedly observed (i.e., microglia polarization and increased density) have often been interpreted as a secondary response to inflammation and neuronal dysfunction ^15,16,46–48^. In line with this hypothesis, it has been demonstrated that microglia (specific genes) were significantly depleted in previously reported hcASD genes^23^. Using a functional burden approach, we demonstrated, however, that variants disrupting microglia-specific genes were in fact associated with ASD, and this was unrelated to known hcASD genes. Although functional burden is not designed to identify exome-wide significant genes, a permutation approach^36^ prioritized 55 genes as (51 novel) candidate contributors to microglia liability for ASD, including *SALL1* and its related network, a well-established central regulator of microglia identity^49^. Among those, we found enrichment for genes involved in the Rho GTPase pathway, which are well-known to be involved in microglia polarization^50,51^. Our findings are consistent with gene expression perturbation studies revealing microglia polarization downstream of hcASD genes that are expressed at low levels in microglia^52^.

Genes identified through genetic association studies and those identified as differentially expressed (DE) in postmortem transcriptomic studies of ASD brains have shown limited (4%) overlap^15,16^. Therefore, ASD-DE genes could not be conclusively interpreted as reflecting primary mechanisms or secondary effects^15^. Using FunBat, we demonstrated that variants disrupting ASD-DE genes significantly contributed to ASD genetic liability. However, we show that ASD-DE genes (both up- and down-regulated) exhibited limited overlap with their corresponding cell-type-specific genes. This is consistent with previous findings showing that the same ASD-DE genes were observed across many cell types^15^. This is in line with schizophrenia^53^, where genetic risk is enriched in DE genes in cerebral cortices of individuals with a diagnosis. We also showed that the cellular liabilities computed with either the cell-type-specific genes or the ASD-DE genes did not converge. Overall, ASD-DE genes may point to broader mechanisms (causes and consequences) that are less cell-type specific.

We observed the well-replicated increase in variants disrupting highly intolerant genes and *de novo* enrichment in females with ASD^41,42^. However, cellular liabilities did not show clear biases towards males or females. Surprisingly, as well, the age of ASD diagnosis did not influence our results on *de novo* enrichment nor on cellular liabilities for ASD. While family trio studies are designed to detect de novo variants, they are less suited for identifying inherited risk alleles since these are highly enriched in the controls (parents and siblings). It is therefore plausible that using unrelated controls may show stronger contributions of inherited risk variants enriched in postnatal and glial cell types.

In conclusion, our study provides a comprehensive analysis of the cellular and developmental mechanisms underlying the genetic contribution to ASD. By identifying novel cell types implicated in ASD and highlighting the variant-class specificity of cellular liability, our findings contribute to a more in-depth understanding of ASD pathology. While all our findings derived from the prefrontal cortex, other cortical areas^37^ and brain structures^14,54^ have been previously associated with ASD, setting the stage for future analyses on the transcriptomic specificities and liabilities of these other brain regions. Future research should build on these findings to develop more targeted and effective interventions for individuals with ASD.

## Methods

### snRNAseq data

To be able to compare effectively the gene expressions across ages, we used single-nuclei RNA sequencing data (snRNAseq) from a unique source [1] with an identical methodology, compared to other available developmental datasets, which aggregate multiple datasets obtained with different technologies[2]. We therefore downloaded and used the 154,748 cells sampled from the prefrontal cortices of 26 individuals aged from gestational week 22 to 40 years old [1]. We used the default quality filters of scanpy (v1.6.0)[3] and AnnData (v0.7.4) as well as the one reported in the original study, to remove i) genes expressed in <5 nuclei; ii) potential nuclei doublets using Scrublet (v0.2.2)[4]; iii) nuclei with gene counts <3 median absolute deviations and <300 genes; iv) nuclei with ribosomal or mitochondrial gene count percentages respectively >20% or >3% per sample MADs ; v) cells with highly expressed MALAT1 gene; and ribosomal and mitochondrial genes. From the 26,747 genes detected, we kept the 15,091 autosomal protein-coding genes. The gene expression within each cell was then scaled to count per million (CPM) and log transformed using scanpy implementation.

We used the 18 cell types defined by the authors using 52 well-established cell-type marker genes[1]. The broad age range was classified into 6 developmental stages[5]: prenatal (gestational week 22—birth); neonatal (birth—2 months); infancy (2 months to 1 year); childhood (1–10 years); adolescence (10–17 years); and adult (17–40 years). We discarded the 1,275 cells previously labeled as poor-quality cells by the authors, as well as the 153 cells for which combinations of cell type X developmental epochs had less than 50 cells to avoid unstable gene assignment. We finally computed the gene specificity for cell types on the 153,325 remaining cells.

### Sensitivity, Specificity, and AUC of Gene Expression Profiles for cell types

We defined the sensitivity of a given gene for a given cell type as the fraction of cells within this cell type that express this gene (read count ≥ 1). The specificity is the fraction of cells outside the cell of interest not expressing the gene. The Receiving Operating Characteristic (ROC) curve represents the trade-off between the sensitivity and specificity through all the possible read count values of a gene (**Figure 1.B**). The Area Under the Curve (AUC) depicted how much the gene has a level of expression unique to the cell type, with values ranging from 0 to 1. A value of 1 indicated that the gene is 100% specific (expressed exclusively in this cell type) and 100% sensitive (expressed in all cells of that cell type). A value of 0.5 indicated that the expression profile of a gene has no discriminatory power for a given cell type (expressed equally in all cells). A value of 0 indicates that the gene is never expressed in the cell type of interest, in contrast to the rest of the cells. The Sensitiity_Max_, Specificity_Max_ and AUC_Max_ correspond to the maximum (best) of these metrics computed across all cell types.

### Defining gene sets, methods to assign genes to cell types

To avoid noisy assignments, we excluded the 7,658 genes with a sensitivity_max_ < 10% (genes detected in fewer than 10% of cells for any given cell type, which we attribute to sparse detection rates in the snRNASeq assay), and found that 86% of the previously defined as “Not detected in the Brain” using bulk transcriptomic data[6] have a sensitivity_max_ inferior to 0.1 (**Supp Figure 1**, **Figure 1.D**).

We compared 4 previously used metrics to assign genes to cell types, including the z-score, the Wilcoxon test, the Expression Proportion (EP), and the Expression Enrichment (EE). For all the methods, *x_i_* represents the average expression of the studied gene in the cluster of interest *i*.

The **z-score** was calculated as follows:

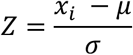

With *μ* the mean of gene expression within the dataset and *σ* the standard deviation:

The **EP** was calculated as follows

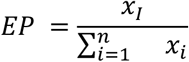

The **EE** was calculated as follows:

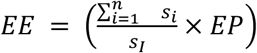

With *s_i_* the total genomic expression in the cluster of interest

The **Wilcoxon test** was computed with the Scanpy implementation. We assigned a fixed number of genes per cell type, as we previously demonstrated that the number of genes assigned impacts burden variance and complexifies interpretation[7]. Finally, to balance both the cell-type specificity and the power for the further burden analysis, we used cell-type gene sets of 600 genes (corresponding to about the top 5% most specific autosomal genes expressed in the cortex). This threshold allows a relatively good number of genetic associations to ASD without saturating the signal (**Figure 2.I**), keeping a good discrimination power of the genes (**Figure 1.E**) and a relatively low overlap between the gene sets (**Figure 1.F**, **Supp figure 3**)

### Validation of the gene sets created with multiple specificity scores

Because the number of genes aggregated for burden association tests influenced the range of effect sizes[7], we created cell-type gene sets of identical numbers of genes to make the cellular liabilities for ASD comparable (**Figure 1.D**). Other studies used up to the top decile of the most specific genes to define each cell type[8]. We therefore wanted to validate that these genes would still be relevant for discriminating between the cell types. For each of the 4 specificity metrics (z-score, EP, EE, Wilcoxon) we selected 15 ranges of 100 genes for each cell type, going from the 100th most specific to the 1,500th most specific. For every set of genes, we computed the PCA based on their expression within the 153,325 cells. The discrimination power was given by the AUC computed on the first component of the PCA[9], showing how reliable the genes are to discriminate the cells of the studied cell type from the rest. While EE discrimination power was close to the z-score we found that EE never assigns 4,116 genes to cell types (at the cell-type specificity threshold of 600 genes). A bootstrap analysis (600 genes randomly sampled 500 times) also showed that the ASD liability only lies in genes not assigned to any cell types (i.e., genes less specific for cell types, as defined by Expression Enrichment (**Supp Figure 17**). Therefore, all the cell-type-specific gene sets used in this analysis originate from the z-score of expression. Finally we showed that genes not under genetic constraint (LOEUF > 0.6) have no liability for ASD (**Supp Figure 5**). We therefore excluded cell-type-specific genes with a LOEUF > 0.6 from the gene sets to prevent dilution of our cellular liability, unless stated otherwise.

### Extraction of gene annotations

The list of genes preferentially expressed in the brain during both prenatal and postnatal stages was extracted from Werling et al.[10] with the prenatal genes corresponding to the *falling* genes and the postnatal genes to the *rising* genes. For the genetic constraint, we extracted the LOEUF (oe.lof.upper) from gnomAD v4.1[11], and for each of the autosomal protein-coding genes, we kept the LOEUF value of the MANE transcript. The ubiquitous and the not-brain-expressed genes were extracted from the HPA (date/version).

### High-confidence ASD genes (hcASD)

The 283 high-confidence ASD genes (hcASD) were defined as the union of 3 previously defined sets of ASD-associated genes: 1) the 232 genes of the SFARI-Gene[12] (Release 2025 Q1) within the category 1 of confidence, 2) the 102 found by Satterstrom et al.[13], with an FDR < 0.05 and the 72 identified by Fu et al.[14] with an FDR < 0.001 as defined in their articles (**Supp figure 1.A**). At the gene contribution step, we used a broader definition of ASD candidate genes by including the 3 categories of ASD genes given by SFARI, as well as the syndromic genes. Moreover, for the genes identified by Fu et al., we included this time the 185 genes with an FDR < 0.05 (**Supp Figure 1.B**).

### Statistical tests

Gene-set enrichments were calculated with a Fisher exact test with a two-sided alternative. The reference used for the enrichment was the 19,152 autosomal protein-coding genes, unless otherwise indicated.

### CNV calling and filtering

DNA was extracted from saliva (OGD-500 kit, DNA Genotek) and genotyped on an Illumina GSA-24v1-0 array (654k SNP sites). We kept only autosomal SNPs with minor allele frequency (MAF) > 5%, probes providing genotypes that are not violating Hardy-Weinberg equilibrium (threshold < 1e-6) and probes with call rates > 90%. Moreover, we used PLINK (1.9)[15] to check for duplicated individuals, sex, and relationships for each participant, and the ones with discordant phenotypic and genetic information were removed. Ancestries (principal components [PC] 1 to 10) were determined with KING[16] (with 3,615 common SNPs), using the standard process defined on the website, and the 1000 Genomes[17] as reference. For the CNV calling, we applied the same methodology as in Huguet et al.[7] (https://github.com/JacquemontLab/MIND-GENESPARALLELCNV) using PennCNV[18] and QuantiSNP[19] algorithms to minimize the type II errors (failure to detect true CNVs) . The following parameters were used for both algorithms: number of consecutive probes for CNV detection >= 3, CNV size >= 1Kb, likelihood scores >= 15. Overlapping CNVs detected by the two algorithms were combined with CNVision[20]. We defined all CNVs with less than 2 copies as deletions and all CNVs with more than 2 copies as duplications. We removed from the following analyses the arrays with an excessive number of detected CNVs (≥ 50), as well as arrays not passing stringent quality-control criteria. We used stringent quality-control criteria: call rate ≥ 95%; standard deviation of log-R-ratio < 0.35; standard deviation of B-allele-frequency < 0.08; and absolute waviness factor < 0.05.

After filtering the arrays based on their quality, we applied filtering for autosomal CNVs. The CNVs with the following criteria were selected for analyses: likelihood score ≥ 30 (maximum of the two CNV calling algorithms), size ≥ 50kb, and overlap with segmental duplications, HLA regions, or centromeric regions < 50%. In addition, we applied an in-house algorithm based on a machine learning method to detect additional artifact CNVs and therefore correct for type I errors (DigCNV, https://github.com/JacquemontLab/DigCNV, details on the training available in Huguet et al.[7]). We annotated the CNVs using GENCODE V19 annotation (hg19) with Ensembl gene name (https://grch37.ensembl.org/index.html). We harmonized CNV frequencies across 4 SPARK batches to prevent spurious associations due to array technology or batch-related differences. For each batch, we computed the frequency of every CNV among the parents to maximize the number of independent individuals and performed a chi-square test to assess frequency heterogeneity across batches (DigCNV, https://github.com/JacquemontLab/DigCNV). The 2,135 unique CNVs showing significant heterogeneity were excluded from further analyses (FDR corrected n=36,360).

### Short variant quality control and annotation

Short variants (i.e., single nucleotide variants (SNVs) and indels) were extracted from individual gVCF files generated for the SPARK iWESv3 cohort[21]. Homozygous reference calls and missing genotypes were excluded, retaining only positions with variant evidence, including SNPs and indels. To increase reliability, only variants detected by both GATK[22] and DeepVariant[23] callers were retained. We then applied numerous thresholds to keep only the most relevant variants (**Supplemental Figure 18**). Genotype-level filtering was then applied using DeepVariant metrics: a minimum read depth (DP) of 20 and a genotype quality (GQ) of 30 were required. The allele count ratio (defined as the ratio of the alternate allele depth over DP) was calculated to assess allelic balance. Because rare variants were not supposed to be homozygous, we removed variants with ratios below 0.2 or above 0.8.

To restrict the dataset to rare variants, we removed any variant with a MAF (Minor Allele Frequency) higher than 1/1,000 individuals for one of these two criteria: (1) the maximum observed allele frequency across all gnomAD v4.1 populations[11], and (2) the allele frequency within the SPARK dataset itself. Functional annotation used Ensembl’s Variant Effect Predictor (VEP[24], release 113.3) aligned to the GRCh38 reference genome. We kept only variants disrupting the MANE transcript definitions[25].

Variants were classified into five categories based on their most severe predicted consequence by VEP (splice variants, frameshift, stop-gained, missense, and synonymous). Splice variants were either splice_acceptor_variant or splice_donor_variant annotations from VEP. Additional filtering criteria were applied per variant class: only LoFtee[11] high-confidence[26] variants were retained for frameshift and stop-gained; only AlphaMissense[27] variants with a pathogenicity score ≥ 0.9 were retained for missense variants, as most of the *likely pathogenic* variants lie above this threshold[27]; and only splice variants with a SpliceAI maximum delta score ≥ 0.8 were included as high-confidence splice variants, as recommended in VEP. The extraction and annotation workflow is available at Zenodo: https://doi.org/10.5281/zenodo.16268986.

### Functional Burden Association Tests (FunBAT)

To estimate the genetic associations of rare variants, we leveraged 4 out of the 6 strategies (strategies 1, 2, 3, and 5) raised in Zuk et al.[28]. We therefore developed in Python an adapted version of previously published models of variant collapsing and summing[7], [29]. For each gene of each individual, we collapsed all variants of the same variant class, resulting in binary information x_i_ (i.e., is the gene disrupted by the variants of interest {1} or not {0}). The functional burden is then computed by summing all the binary information for all the genes of the gene set of interest for each individual (Formula 1).

Because deletions and duplications can be multigenic, a simple burden model would not inform on the effect of a given gene but on the cumulative effect of all the disrupted genes. We therefore adapted the model to counterbalance the effect of the other genes impacted by the multigenic variant but not included in the studied functional gene set. We added the average effect of the rest of the genome by computing two burdens, one for the genes under genetic constraint (LOEUF < 1) and a second one for genes not under genetic constraint (LOEUF > 1) (Formula 2).

The ASD liability of the studied function corresponds to the β1 of the following logistic regression. Because individuals in the dataset were not independent, we used a conditional logistic regression model to correct for data relatedness by using the ConditionalLogit function implemented in the statsmodels Python library. As diversity within the groups is a prerequisite for conditional regression analysis, we removed from the analysis the 12,955 groups (families) of single individuals or with only ASD diagnoses, corresponding to a total of 16,910 individuals. The final numbers of individuals used for the models are described in **Supp Table 1**. Regression models are also corrected for the sex of the individuals as well as their ancestry through the use of the first 10 PCs given by PLINK[15] and the number of rare synonymous variants for each individual.

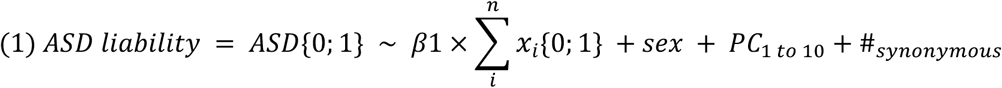

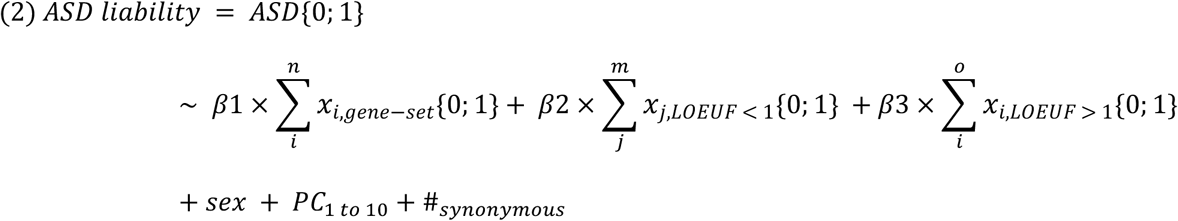

A standalone version of the FunBAT allowing users to compute their functional burden association tests is available through the Python package FunBAT (https://github.com/JacquemontLab/FunBAT).

We asked whether varying the levels of cell type specificity of our gene sets influenced the findings described above. To do so, we recomputed burden tests at increasing levels of specificity from the top 90th (n∼1,000 genes) to the top 99th percentile (n∼100 genes) of gene specificity for each cell type (**Figure 2.I**). We also asked if varying the sample size of our genetic population influenced our findings. We therefore recomputed the cellular liabilities after randomly sampling families from our datasets. We tested 9 sizes of downsampling (from 10% to 90% with a 10% step). We computed the cellular liabilities for ASD following the same process as in figure 2.G. This process was repeated 20 times, and then we computed the correlations of the downsampled liabilities to the original one with Spearman correlations.

### *De novo* analysis

We identified the *de novo* variants in 31,814 offspring from 20,130 families. A variant was defined as *de novo* if neither of the two parents had the variant identified in one read or more, regardless of the variant quality control. Following previous studies[30], we kept only one *de novo* variant per gene by selecting the one with the strongest consequence (as defined by Ensembl[24]). We removed 28 individuals from the analyses with abnormally high numbers of *de novo* synonymous alterations (> 10 genes with *de novo* synonymous). We computed *de novo* enrichments for cortically expressed genes under constraint (LOEUF < 0.6[11]). Consistent with previous studies on the SPARK dataset[30], we found that 28% of non-ASD individuals have at least one *de novo* synonymous variant. The rate of *de novo* synonymous variants among non-ASD females and males was similar (Fisher exact test, p-value = 0.5). The enrichments of *de novo* variants across the cell-type-specific gene sets were computed using a Fisher’s exact test. In the first analysis, we used a contingency table reporting the number of *de novo* and inherited variants in a given gene set compared to the de novo and inherited variants across all cortically expressed genes. For the second, we created a contingency table for each gene set, reporting the number of de novo and inherited variants in ASD individuals compared to the de novo and inherited variants in non-ASD siblings. While not depicting the same information (the enrichment of de novo variants compared to the other cell types on one hand and the enrichment of de novo variants in ASD compared to non-ASD for a given cell type on the other hand), the enrichments computed with the two methods were correlated for all variants (rs = 0.31 to 0.60, p-value = 2e-3 to 3e-9) outside splice variants.

### Gene-set subsampling analysis

We adapted a previously published method[31] to prioritize individual genes contributing to the significant cell-type functional burden tests. We developed a bootstrap version of FunBAT to test the association between downsampled gene sets and ASD. For each, we selected the developmental epochs in which we observed significant association with ASD when the hcASD genes are removed. For each cell type, we generated 1,000 sub-gene sets by randomly sampling 100 genes from the initial cell type-specific gene set. We then recomputed the ASD burden for these 1000 sub-gene sets following the same method as for the initial gene sets and identified the sub-gene sets significantly associated with ASD (FDR < 0.05, n = 1,000). Genes were prioritized if they were enriched among the significantly associated sub-gene sets (Fisher’s exact test, FDR < 0.05).

### Comparing variance and burden approaches

To identify whether the ASD liability of cell types stemmed from a small portion of its specific genes or resulted from the effect of a large proportion of its genes, we compared the p-value of two methods implemented in the R-package SKAT[32]. We applied the SKAT algorithm on the collapsed information of genes for each variant type. The SKAT models (both burden and variance) were corrected by the sex and the ten first principal components, like our models. Of note, SKAT-burden provides identical results to FunBAT when using classical logistic regression and slightly better results than the conditional logistic regression model (**Supp Figure 19**). This can be explained by the loss of individuals who are part of families without ASD diversity (i.e., families of only one individual or families with all the members diagnosed with ASD).

### Analysis of the ASD-DEG

We extracted the ASD-DE genes identified in the 36 cell types from Wamsley et al.[33], using the same threshold (FDR < 0.05, log2FC > 0.2) as the authors. We kept only the 47 sets of ASD-DE genes containing more than 100 DE genes. Because the sets of DEG genes were of different sizes and the burden method is influenced by the gene-set size (**Supp Figure 20**)[7], we can’t directly compare their effect sizes on ASD. We therefore computed the distribution of effect sizes for 540 random gene sets of different sizes, ranging from 50 to 950 with a step of 50 (30 gene sets per size). The normalized effect sizes of the ASD-DE gene sets were based on the mean and SD computed on the 30 sets of closest size. For each cell type, we wanted to compare the liabilities of the ASD-DE gene sets and the ones of the cell-type-specific gene sets. We extracted from the cell x gene expression matrix all the cells coming from control individuals and then applied the same method as the one used for the Herring et al. dataset (c.f. Methods). The functional burden was computed for the 36 cell-type-specific gene sets across the same 6 gene-disrupting variant classes. We validated the consistency of our findings by correlating the effect sizes computed for these cell types and their equivalent ones from the Herring (**Supp Figure 9.B**).

### Analysis of liability biases computed between ASD population subsets

As ASD is a spectrum, we tried to identify different cellular liability for ASD when different subtypes of ASD were studied. We therefore tested whether different cell types or developmental epochs were in action for male ASD compared to female ASD. We recomputed the functional burden of the 91 cell types for these different ASD subgroups using the same method as before. To identify the significant difference between the two groups tested, we performed a permutation test where the ASD individuals were randomly split 5,000 times following the same proportion as the sex ratio in the ASD dataset (i.e., 0.27 for female and 0.73 for male). Biases in cellular liabilities (differences of OR) were defined as significant when the difference was smaller or larger than 5% and 95% of the differences computed for the 5,000 permutations. These empirical p-values were hence corrected for multitesting (FDR). For the age of diagnosis, we split the dataset again into two: the early diagnosis for ASD individuals diagnosed before the age of 4 (50% of ASD) and the late diagnosis for the ASD individuals diagnosed later than 4 (50% of ASD). We hence repeated the same method used for the sex bias analysis across the age of diagnosis. Because males and females were not diagnosed at the same age (i.e., medians = 62 and 44 months, Mann-Whitney U-test p-value < 1e-300, **Supp Figure 21**), we tested *de novo* excesses between age groups for the two sexes separately.

### Resource availability

Lead contact: For additional information, as well as requests regarding resources, please direct your inquiries to the lead contact:

Renne T.

Jacquemont S.

## Data availability

SPARK population data (both genetic and phenotypic) are available to other investigators online (https://base.sfari.org/). Derived measures at the individual level used in this study are available upon request (S.J.). Summary statistics and the gene sets used to compute them have been deposited on Zenodo: https://doi.org/10.5281/zenodo.17822332.

## Code availability

All original scripts have been deposited and are publicly available as of the date of publication on GitHub or Zenodo repositories: 1) Quality control and annotation of short variants : https://doi.org/10.5281/zenodo.16268986; 2) Quality control and annotation of CNVs : https://github.com/JacquemontLab/MIND-GENESPARALLELCNV and https://github.com/JacquemontLab/DigCNV; 3) FunBAT algorithm: https://github.com/JacquemontLab/FunBAT; 4) Figures and other scripts used to create gene sets and analyze FunBAT results: https://doi.org/10.5281/zenodo.17860027.

## Acknowledgments

This research was enabled by support provided by Calcul Quebec (http://www.calculquebec.ca) and the Digital Research Alliance of Canada (https://www.alliancecan.ca/). Sebastien Jacquemont is a recipient of a Canada Research Chair in neurodevelopmental disorders and a chair from the Jeanne et Jean Louis Levesque Foundation. This work is supported by a grant from the NIH: 5U01MH119690. We are grateful to all of the families in SPARK, the SPARK clinical sites and SPARK staff. We appreciate obtaining access to phenotypic and genetic data on SFARI Base. Approved researchers can obtain the SPARK population dataset described in this study by applying at https://base.sfari.org. We appreciate obtaining access to recruit participants through SPARK research match on SFARI Base.

## Author contribution

Conceptualization: T.R., G.H., T.N., S.J.; Data curation: T.R., F.B., V.-R.B., G.H.; Formal Analysis: T.R., A.D.; Funding acquisition: S.J.; Investigation: T.R., A.D.; Methodology: T.R., F.B., C.P., G.H.; Project administration: G.H., S.J.; Resources: T.R., F.B., C.P.; Software: T.R., F.B., C.P.; Supervision: T.N., S.J., G.H.; Validation: T.R., S.J.; Visualization: T.R., S.J.; Writing – original draft: T.R., T.N., S.J.; Writing – review & editing: T.R., F.B., C.P., A.D., V.-R.B.; G.H., T.N., S.J.;

