## Supplementary Figures for "Genetic and Cortical Cell-Type Liability Architecture of Autism"

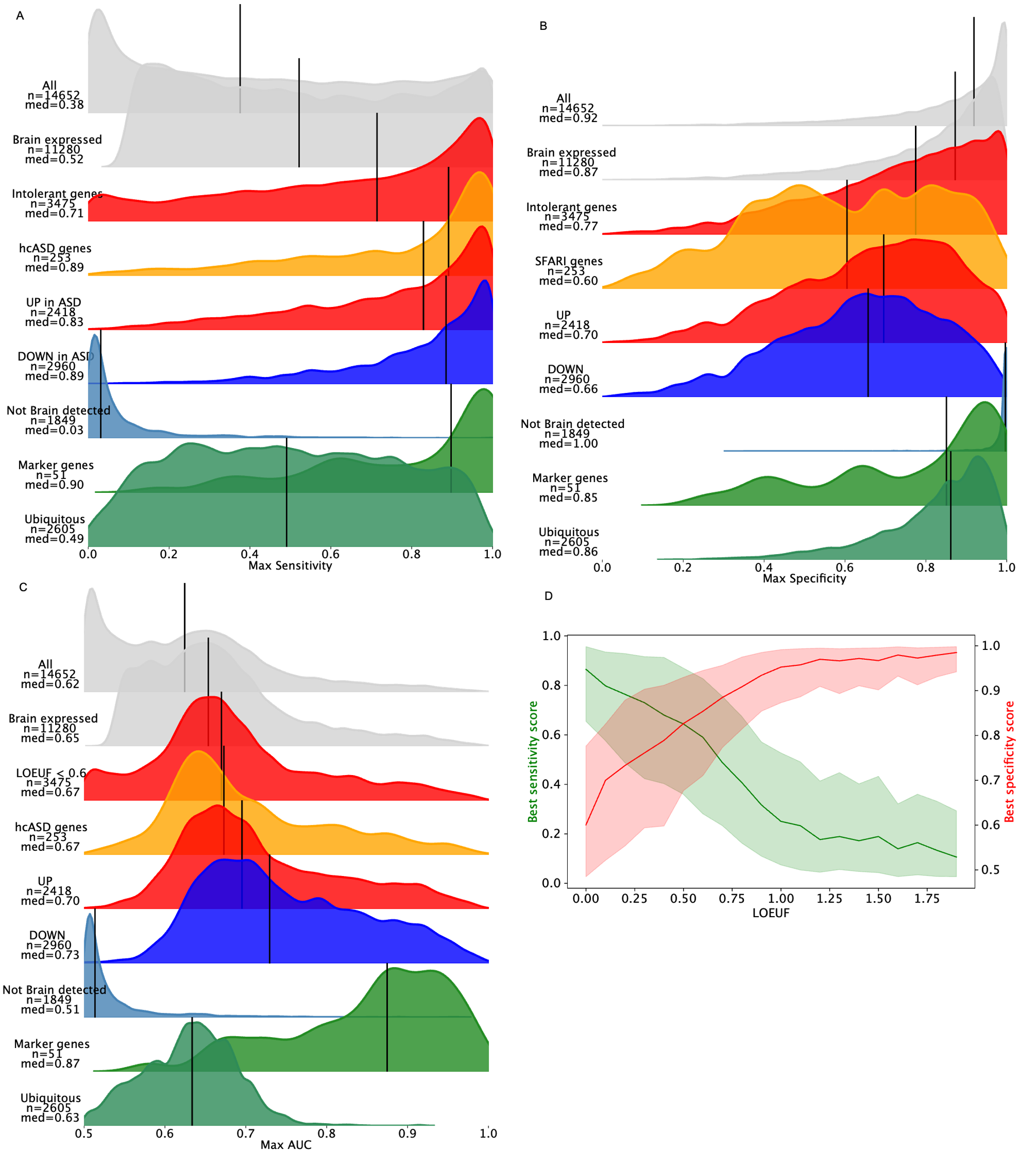
**Supplementary Figure 1**: **Distributions of sensitivity and specificity of multiple gene sets.** Distribution of the maximum sensitivity (A), specificity (B) and the resulting AUC (C) distribution of the maximum specificity computed across the 91 developmental cell types identified in the snRNAseq dataset[1]. Medians are depicted by black lines. Brain expressed genes were by definition all the protein-coding genes with a sensitivity > 0.1. Intolerant genes correspond to the protein-coding genes with a LOEUF v4.1[2] < 0.6 for their MANE transcript. Genes not detected in the brain and the ubiquitous (genes with a low cell type specificity) were extracted from the human proteome atlas (<https://www.proteinatlas.org/>). The marker genes are the one used to create the 18 cell types of the dataset[1].

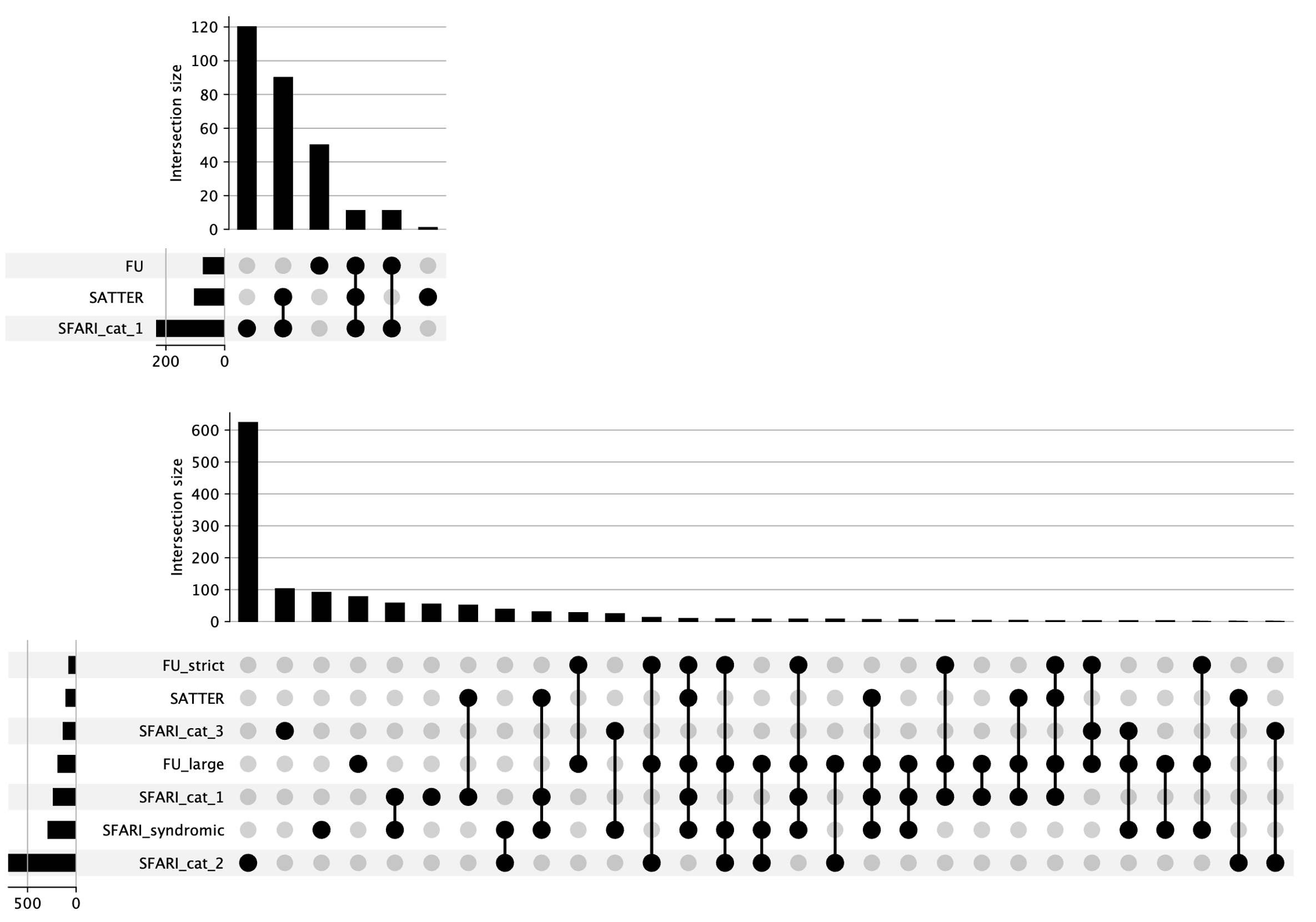

**Supplementary Figure 2:** **Distribution of the high-confidence ASD genes.** Upset plot representing where the 283 hcASD genes are coming from as well as the overlap between the 3 sources[3], [4], [5]. The horizontal bar chart represents the size of the three used sets of ASD-associated genes. The vertical bar chart indicates the number of genes in each unique or shared set, while the filled circles below represent the combinations of datasets contributing to each intersection.

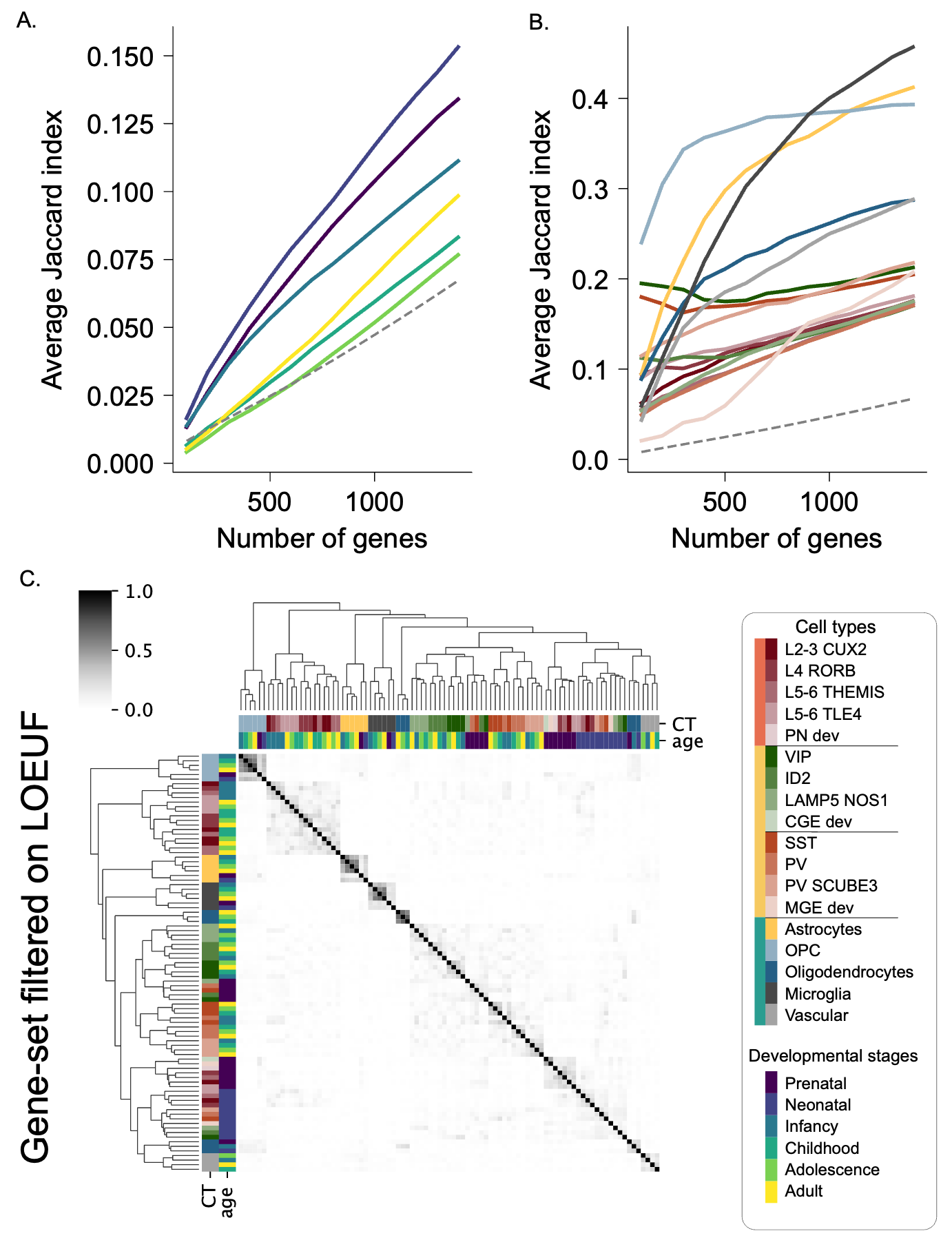

**Supp Figure 3:** **Matrix of Jaccard indexes between the cell type gene sets**. Jaccard indexes in a gray color scale are computed between each of the 91 developmental cell types. Cell types are clustered based on the matrix of Jaccard indexes with the Ward method. The dendrograms on the axes show the hierarchical clustering of the gene sets based on their overlapping sets of genes.

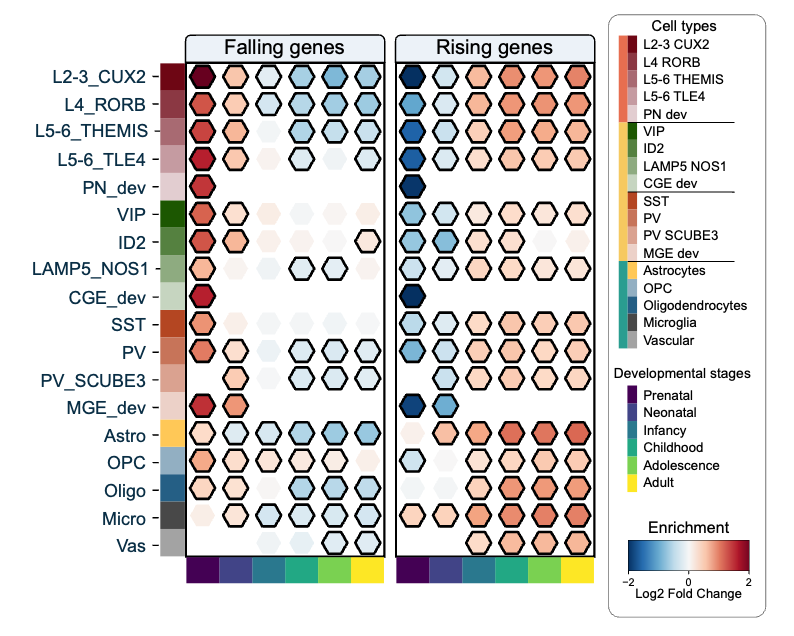

**Supp Figure 4: Matrix of enrichment for pre- and postnatal genes.** Matrix representing the enrichment of the 91 cell types (18 cell types in rows and 6 developmental epochs in columns) for the falling and rising genes[6]. Falling genes are defined as genes with higher expression during the prenatal stage, while rising genes have higher expression during the postnatal stage. The color scale indicates the Log2 Fold Change enrichment, ranging from -2 to 2.

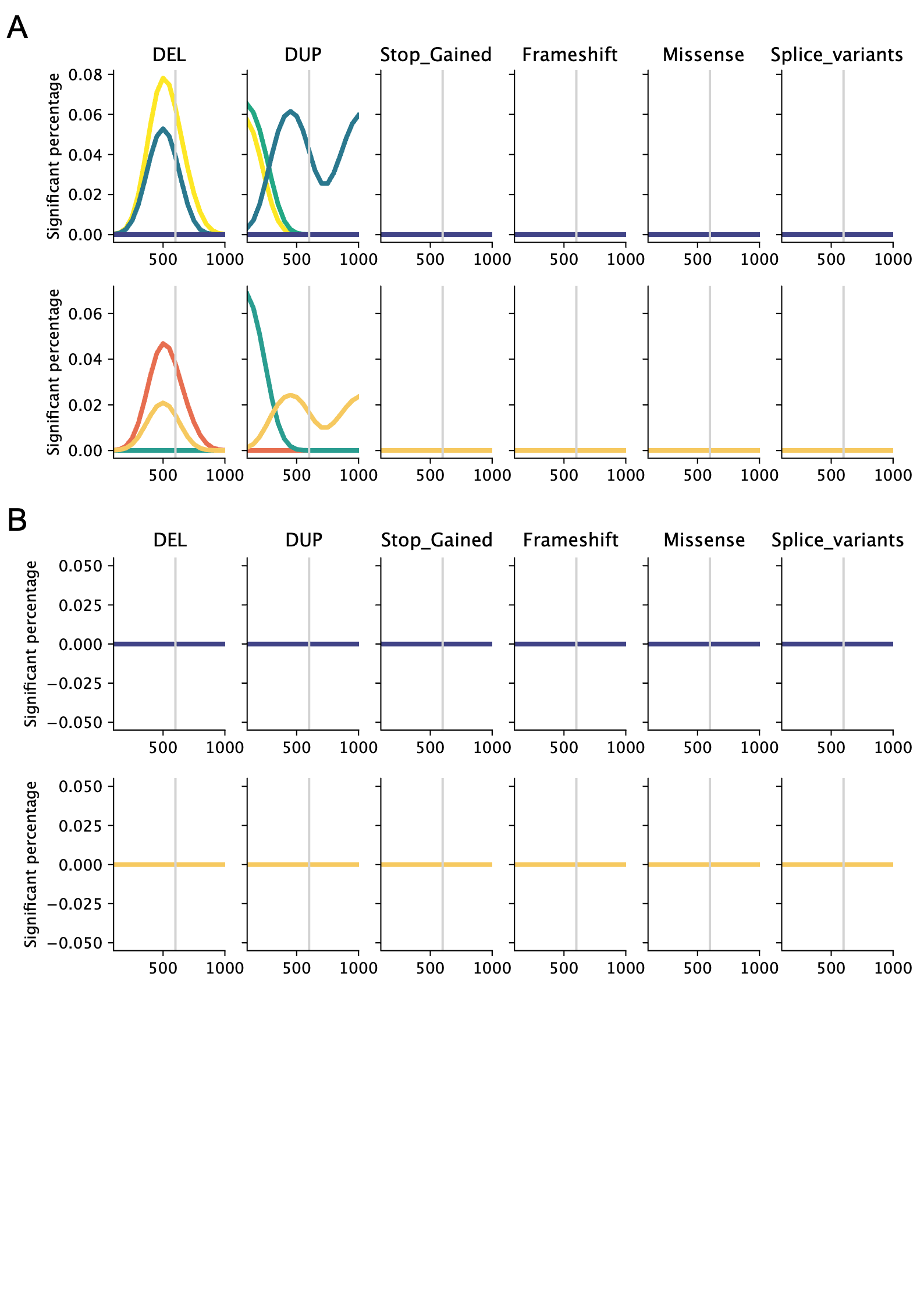

**Supplementary Figure 5:** **Percentage of cell types with a significant liability for ASD when their genes not under genetic constraint[2] are disrupted.** The analysis is split into two (A) for genes moderately intolerant (LOEUF between 0.6 and 1) and (B) for tolerant genes (LOEUF > 1), The Y-axis represents the percentage of cell types significantly associated with ASD when grouped on the main cell classes (top) and the developmental stages (bottom). The X-axis represents the size of the gene sets ranging from the top 99th (n=100) to the top 90th (n=1100) percentile of genes with the highest relative expression in each cell type. The different types of rare gene-disrupting mutations are shown in separate plots.

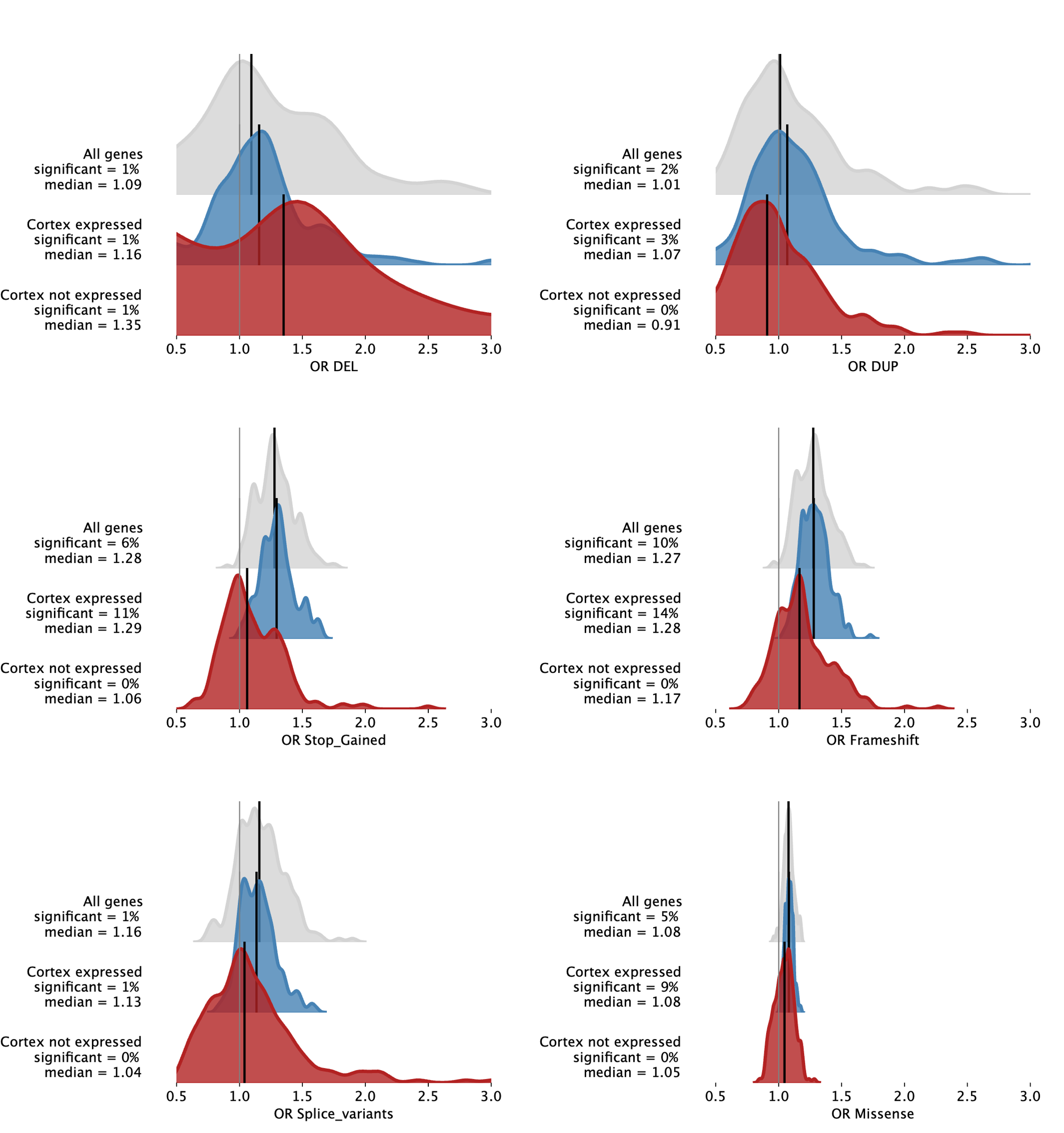
**Supp Figure 6:  Distribution of ASD liability for 2 genes expressed in the cortex or not.** The figure shows the distribution of genetic liabilities for Autism Spectrum Disorder (ASD) for six types of gene-disrupting variants: DEL, DUP, Stop_Gained, Frameshift, Missense, and Splice_variants. The distributions are shown for three categories of genes: all protein-coding genes (in grey), genes identified in the prefrontal cortex (in blue), and genes not observed in the prefrontal cortex (never observed in more than 10% of a cell type, in red). Each plot displays the significance percentage and median value for the genetic liability of ASD for each category.

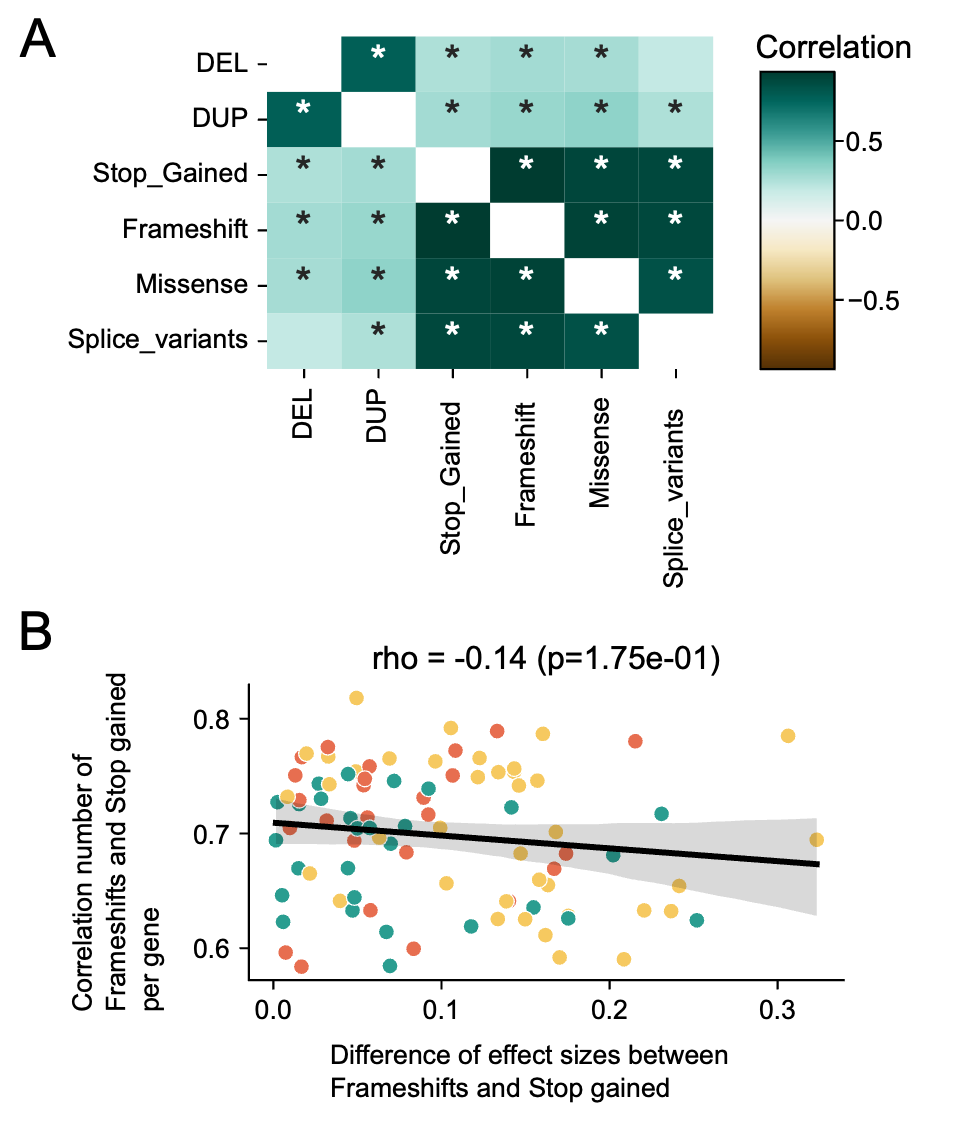

**Supplementary Figure 7: Comparison of the number of variants per gene sets and per gene.** (A) Matrix representing pairwise correlations between the number of variants disrupting genes of the 91 cell types. The color scale represents the Spearman r-value, and stars depict significant correlations. (B) Scatterplot representing the absence of a link between the divergence of ASD liabilities computed for stop gained and frame shifts (X-axis) and the correlation of the number of variants per gene for each cell type (Y-axis). The 91 cell types are colored based on the 3 main cell classes (green for glia, orange for excitatory neurons, and yellow for inhibitory neurons).

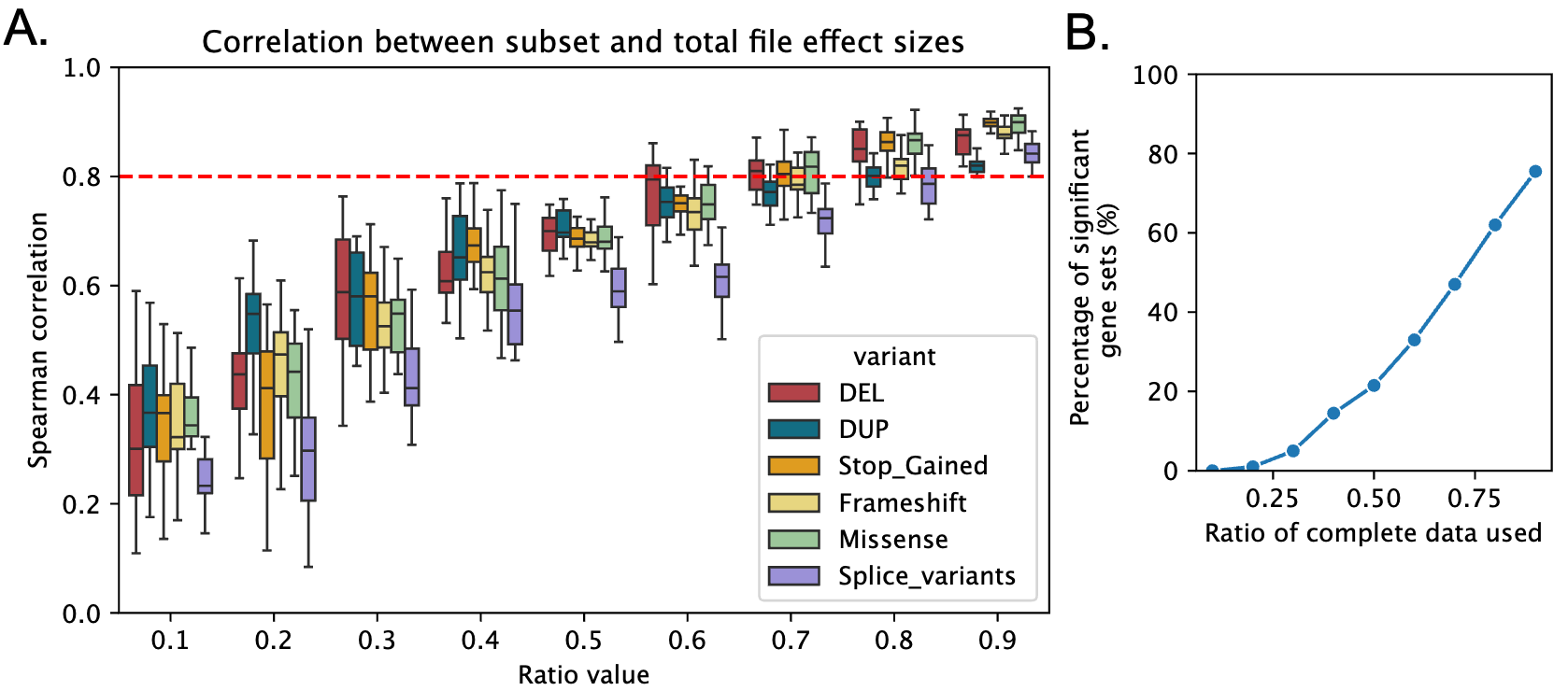

**Supplementary Figure 8. Sensitivity analysis of the cellular liability for ASD along the population size.** (A) Spearman's correlation between the cellular liabilities computed on the total population (Figure 2.G) and randomly sampled individuals from the total population of 10 different sizes (20 samples per size, going from 10% to 90% of the initial dataset). The correlations are computed for liabilities computed for each variant type individually. The red dotted line represents the threshold for highly correlated results. (B) Subanalysis of the microglia sets significantly associated with ASD when disrupted by stop gained and frameshift. The plot line represents the percentage of the cell types that were also significantly associated with ASD when analyzing only a subsample of the initial dataset.

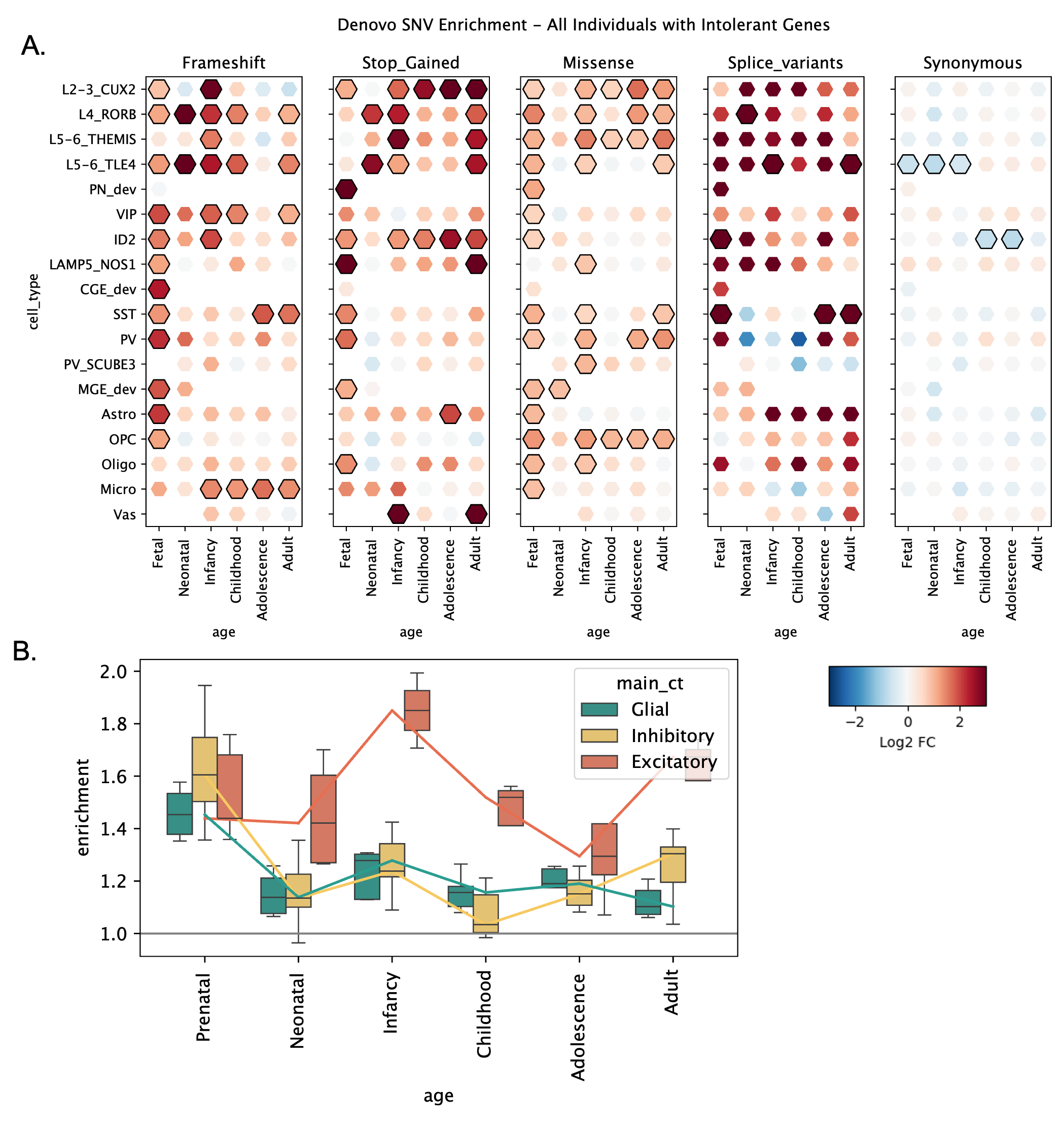

**Supplementary Figure 9: Enrichment of de novo variants within the ASD individuals compared to unaffected siblings.** (A) Enrichment of de novo variant rate for the 91 cell types x developmental stages and 5 variant classes. Enrichments correspond to the rate of de novo/inherited variants for all the under-constraint (LOEUF < 0.6) genes assigned to each developmental cell type between the ASD individuals and the unaffected siblings. The log2FoldChange of the enrichment is color-coded with a scale going from depleted (blue) to enriched (red). Significant enrichments are outlined in black. (B) Distribution of the enrichments (fold change) of de novo variants grouped in the 3 main cell classes and 6 developmental times for the 4 types of gene-disrupting variants (stop gained, frameshift, splice variants, and missense variants). Values above 1 represent cell types enriched in de novo variants relative to the unaffected siblings.

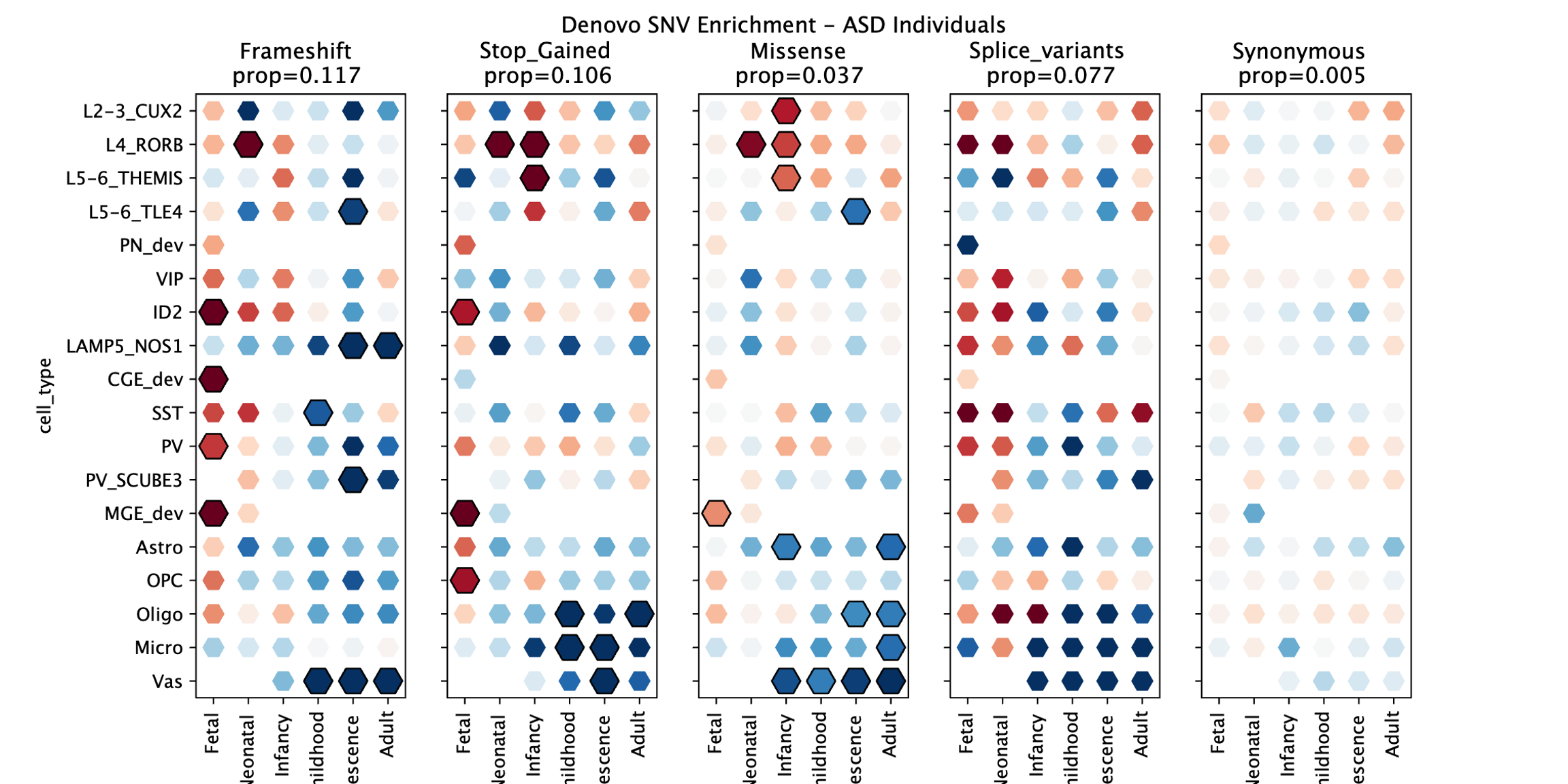
 **Supplementary Figure 10: For ASD individuals, enrichment of de novo variants within each cell type compared to the rest of brain-expressed genes.** (A) Enrichment of de novo variant rate for the 91 cell types x developmental stages and 5 variant classes. Enrichments correspond to the rate of de novo/inherited variants for all the under-constraint (LOEUF < 0.6) genes assigned to each developmental cell type between the ASD individuals and the rate of all other brain-expressed genes (also under genetic-constraint). The log2FoldChange of the enrichment is color-coded with a scale going from depleted (blue) to enriched (red). Significant enrichments are outlined in black.

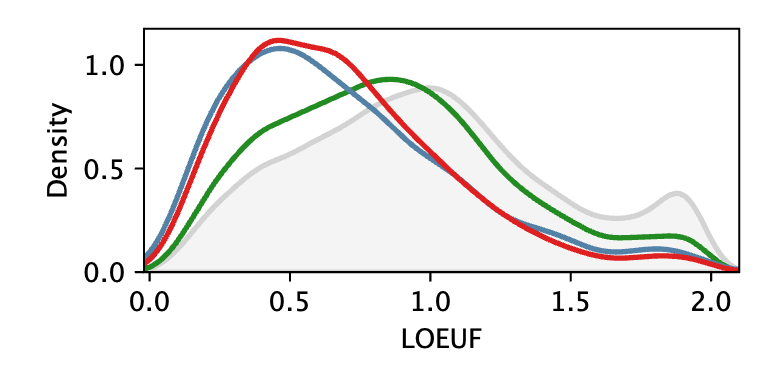

**Supplementary Figure 11: LOEUF distribution of the genes assigned to the 36 cortical cell types in Wamsley et al..** The ASD upregulated genes, the ASD downregulated genes, and the cell type specific genes identified in Wamsley et al are represented in blue, red, and green, respectively.

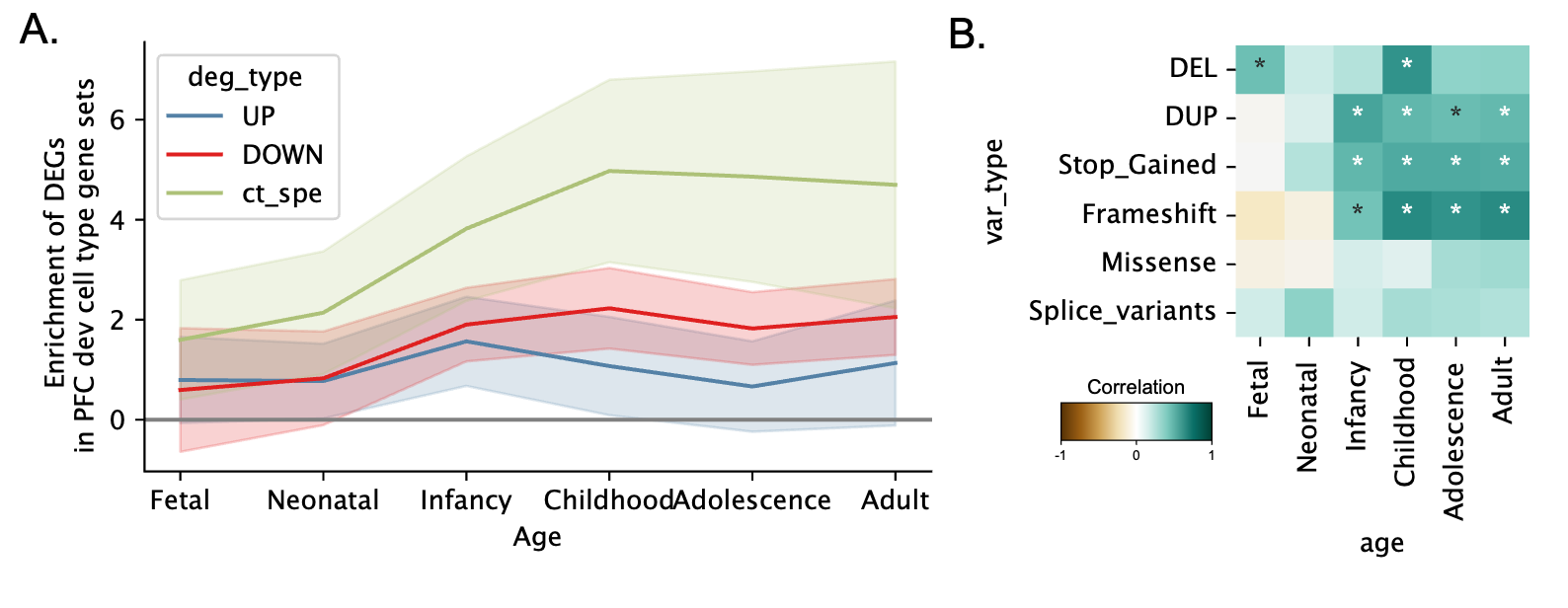

**Supplementary Figure 12: Comparison between Wamsley cell type gene sets and Herring gene sets.** (A) Enrichment of the gene sets computed from the 3 gene assignment methods (upregulated in blue, downregulated in red, and cell type specific in green) with their corresponding Herring cell types (**Supp. table 5**) at 6 developmental timepoints. (B) Correlation of the ASD liabilities computed for each class of variants between the Wamsley cell type specific gene sets and the Herring cell type specific gene sets, at the 6 developmental stages. The color code represents the level of Spearman correlations, and stars represent the significant ones.

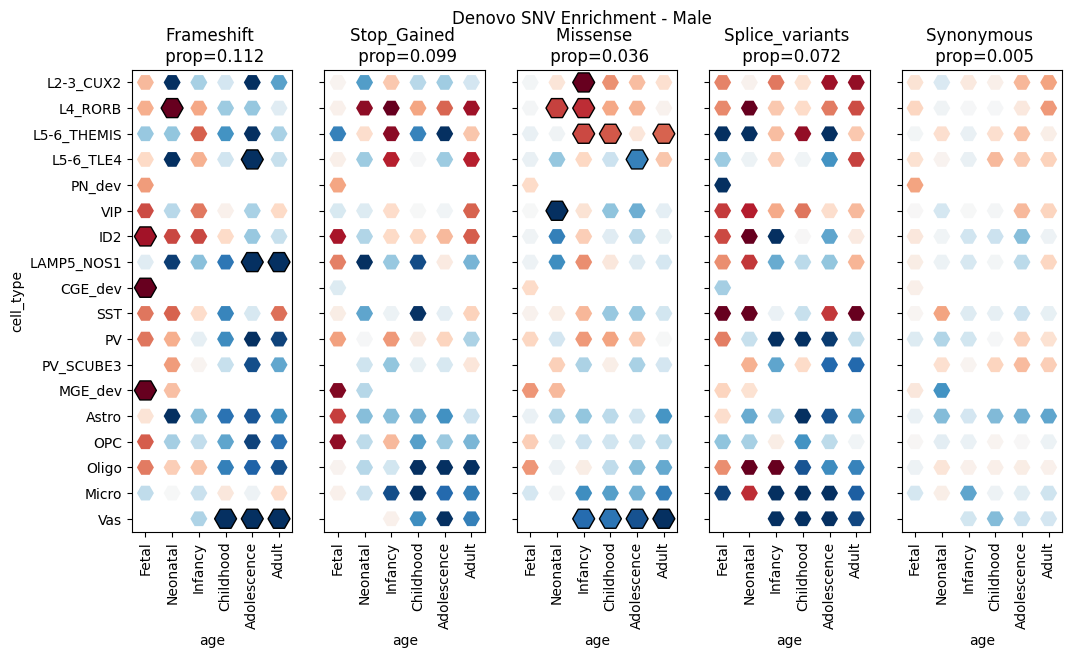

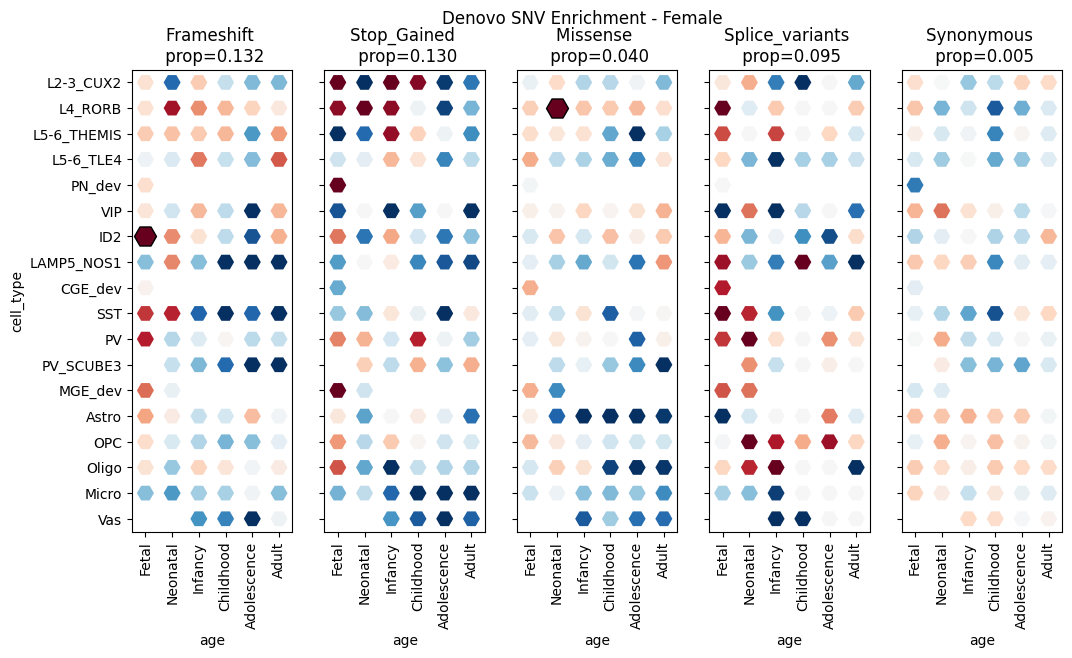

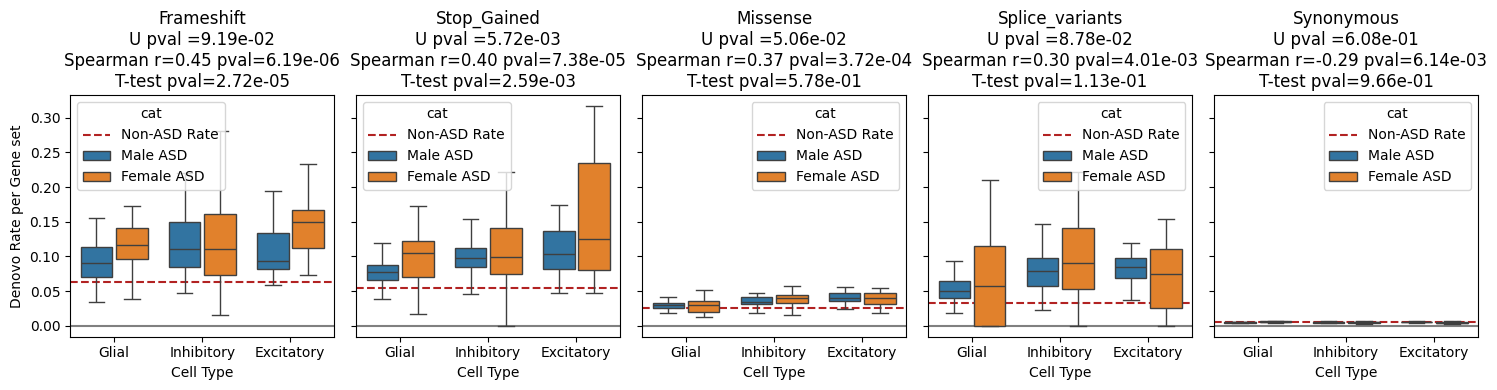
**Supplementary Figure 13. Denovo analysis in cell type gene sets for the ASD males and ASD females.** Distribution of the de novo enrichment computed for the genes under constraint (LOEUF < 0.6) across the 91 cell types for the ASD males (A) and ASD females (B), respectively. (C) Comparison of the de novo enrichment computed between the two ASD subsets for the 91 cell types clustered into 3 categories following the 3 main cell classes (i.e., excitatory neurons, inhibitory neurons, and glia).

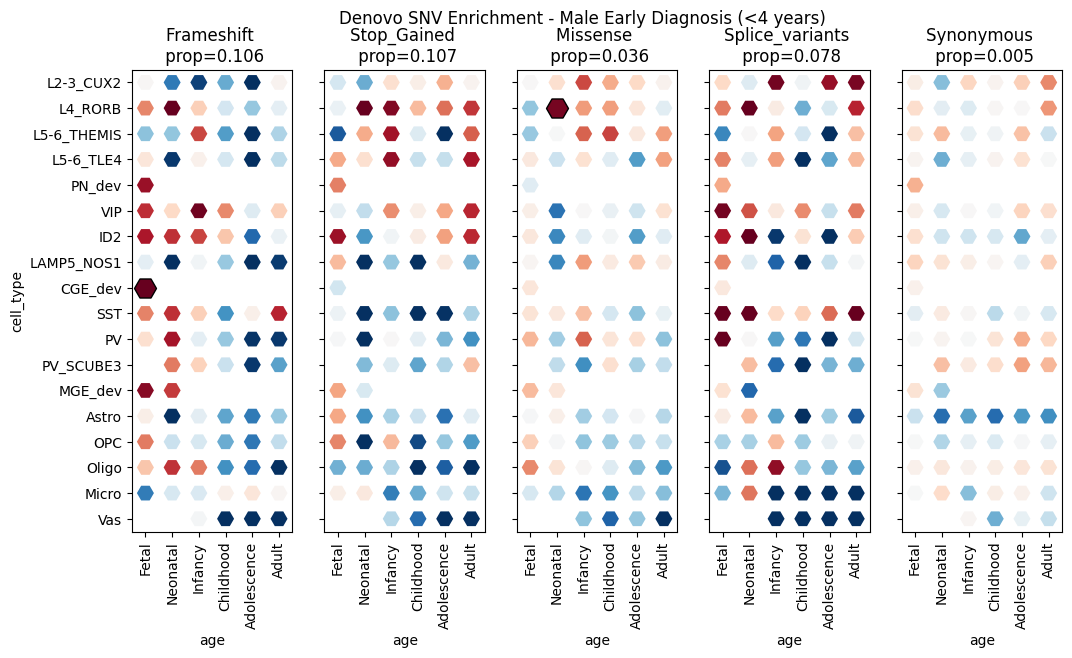

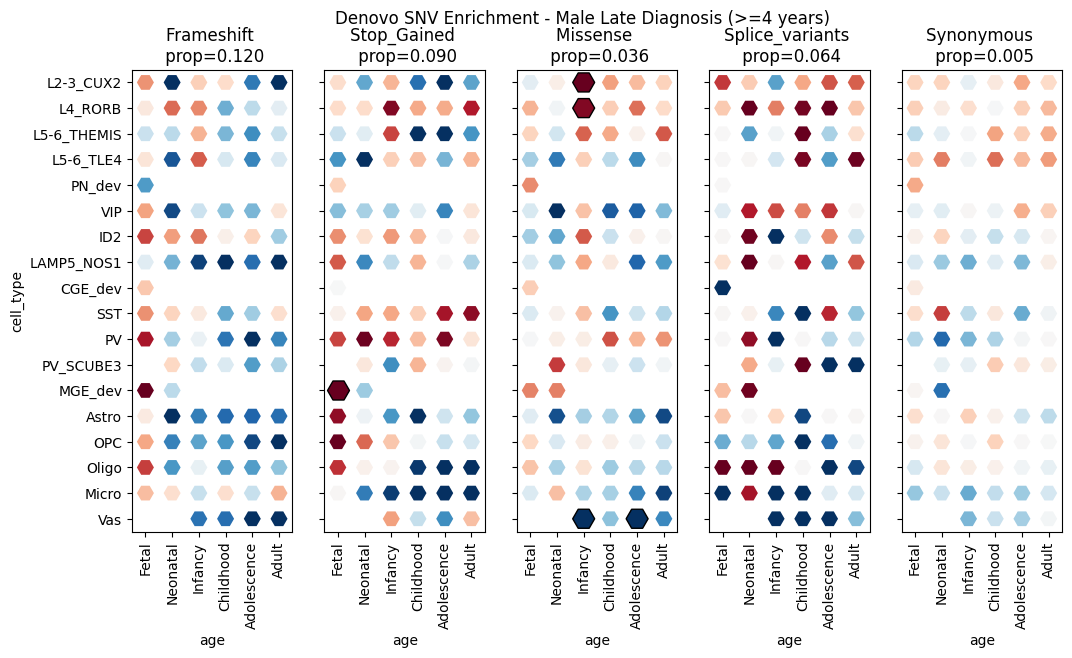

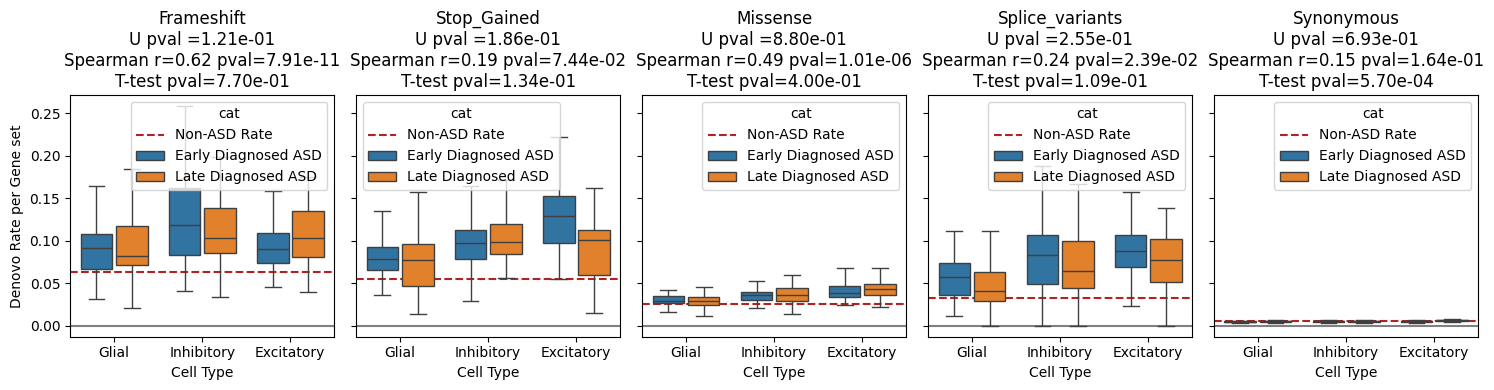

**Supplementary Figure 14. *De novo* analysis in cell type gene sets for early-diagnosed ASD and late-diagnosed ASD in males.** Distribution of the *de novo* enrichment computed for the genes under constraint (LOEUF < 0.6) across the 91 cell types for (A) the early diagnosed ASD (before 4 years old) and (B) the late diagnosed ASD (after 4 years old), respectively. (C) Comparison of the *de novo* enrichment computed between the two ASD subsets for the 91 cell types clustered into 3 categories following the 3 main cell classes (i.e., excitatory neurons, inhibitory neurons, and glia).

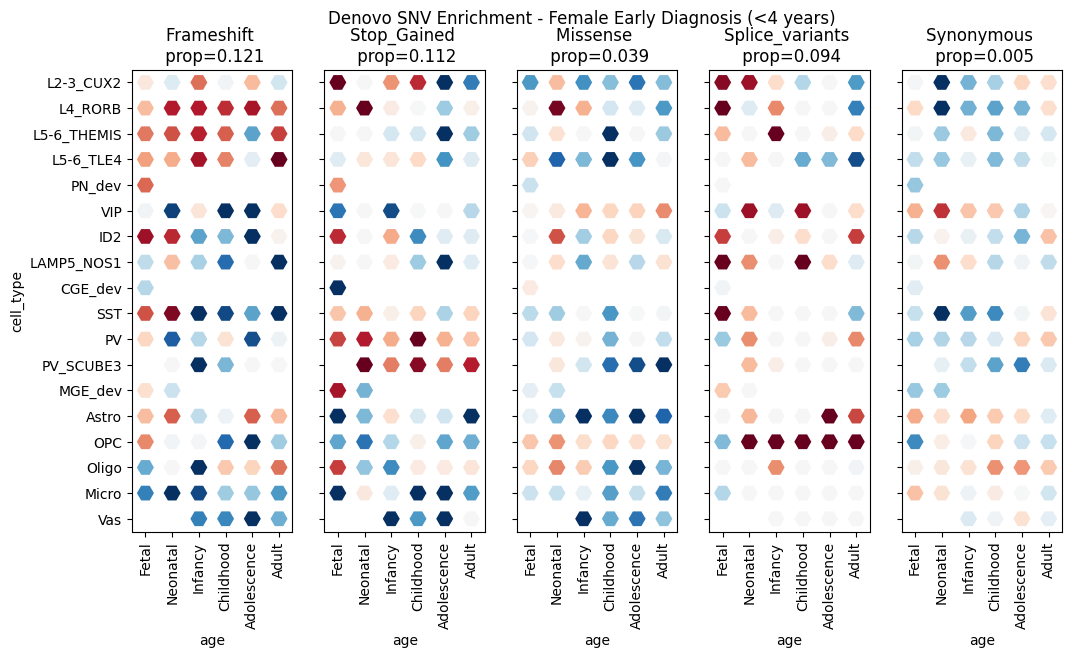

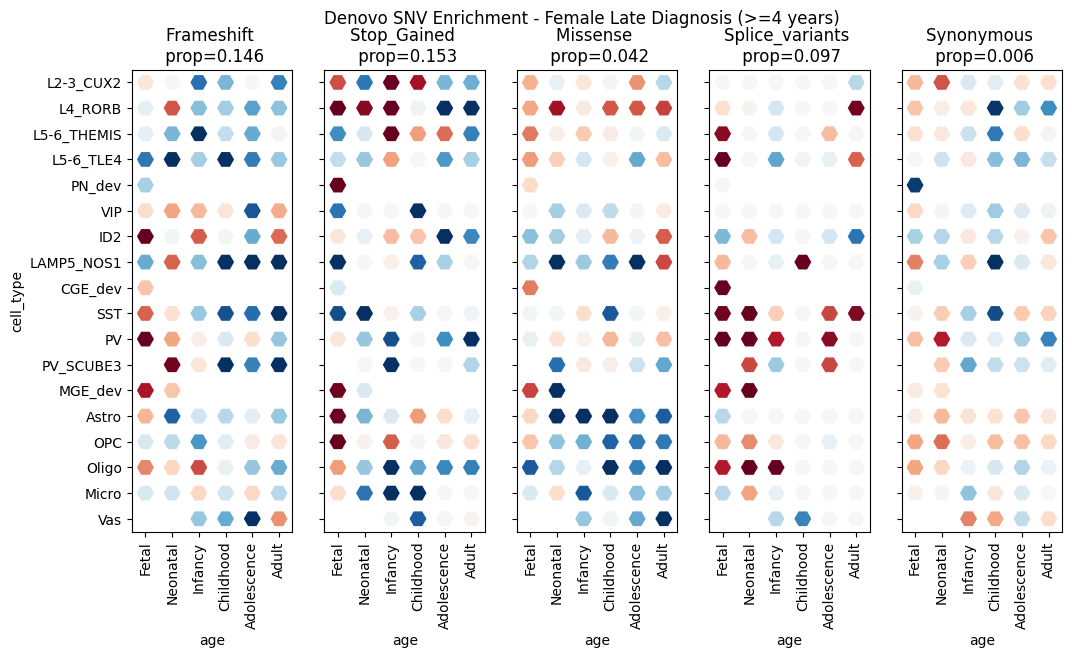

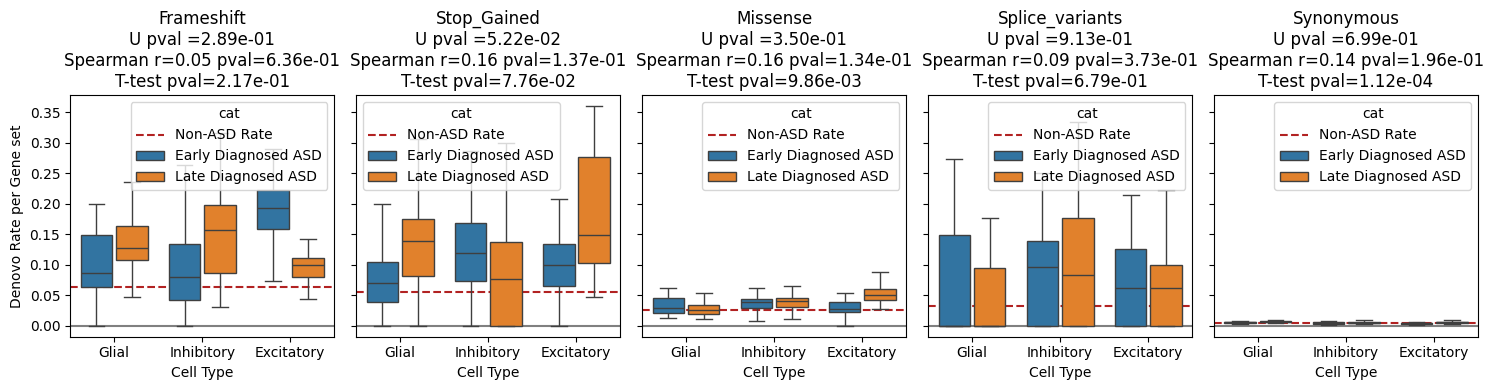

**Supplementary Figure 15. *De novo* analysis in cell type gene sets for early-diagnosed ASD and late-diagnosed ASD in females.** Distribution of the *de novo* enrichment computed for the genes under constraint (LOEUF < 0.6) across the 91 cell types for (A) the early diagnosed ASD (before 4 years old) and (B) the late diagnosed ASD (after 4 years old), respectively. (C) Comparison of the *de novo* enrichment computed between the two ASD subsets for the 91 cell types clustered into 3 categories following the 3 main cell classes (i.e., excitatory neurons, inhibitory neurons, and glia).

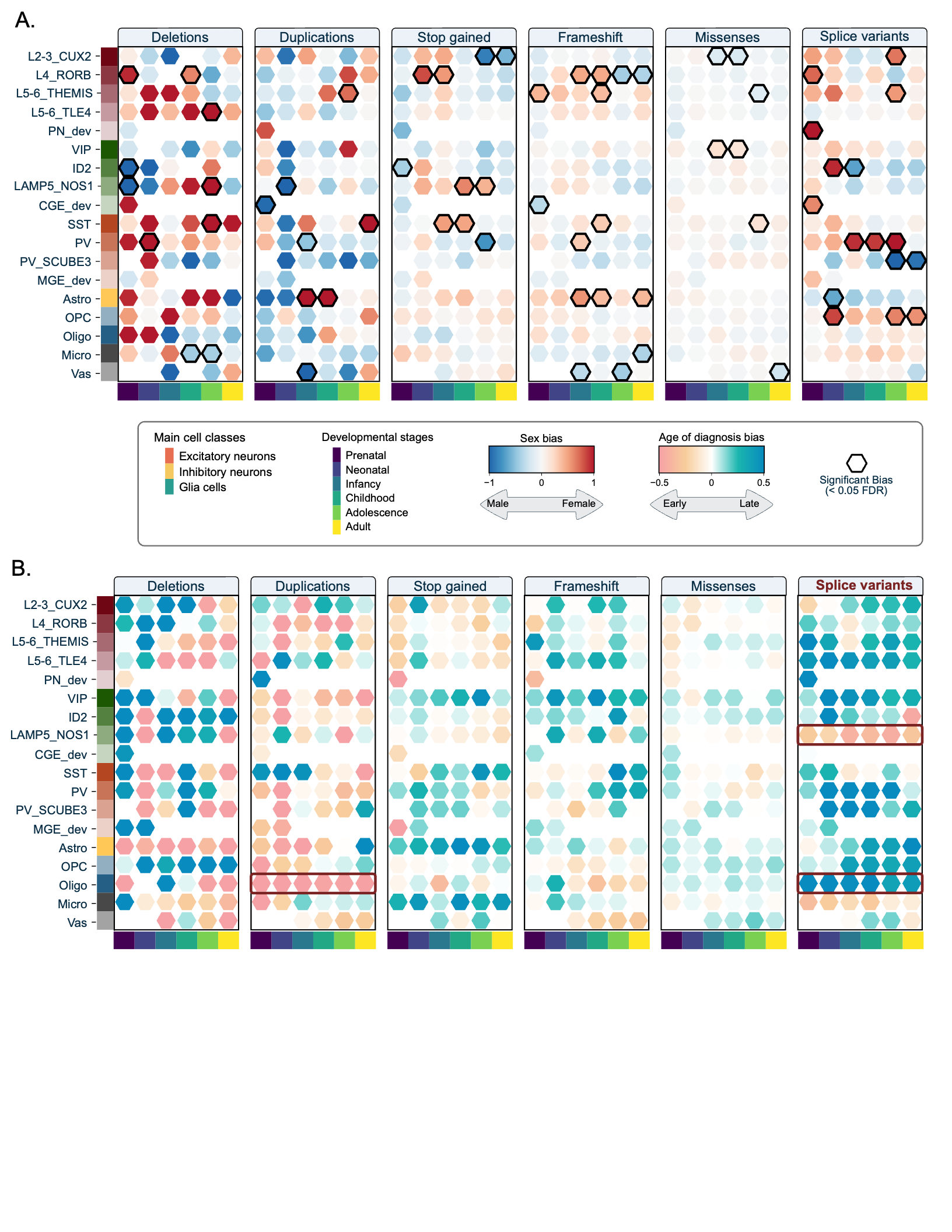

**Supplementary Figure 16. Cellular liability biases computed for different subsets of ASD.** (A) Sex bias of the cellular liabilities computed for the 91 developmental cell types across 6 types of variants. The bias is color-coding the bias toward males (blue) and females (red). Significant biases (empirical p-value < 0.05) are outlined in black. (B) Age-at-diagnosis bias of the cellular liabilities computed for the 91 developmental cell types across 6 types of variants. The bias is color-coding the bias toward earlier diagnosis (orange) and later diagnosis (blue). Significant biases (empirical p-value < 0.05) are outlined in black.

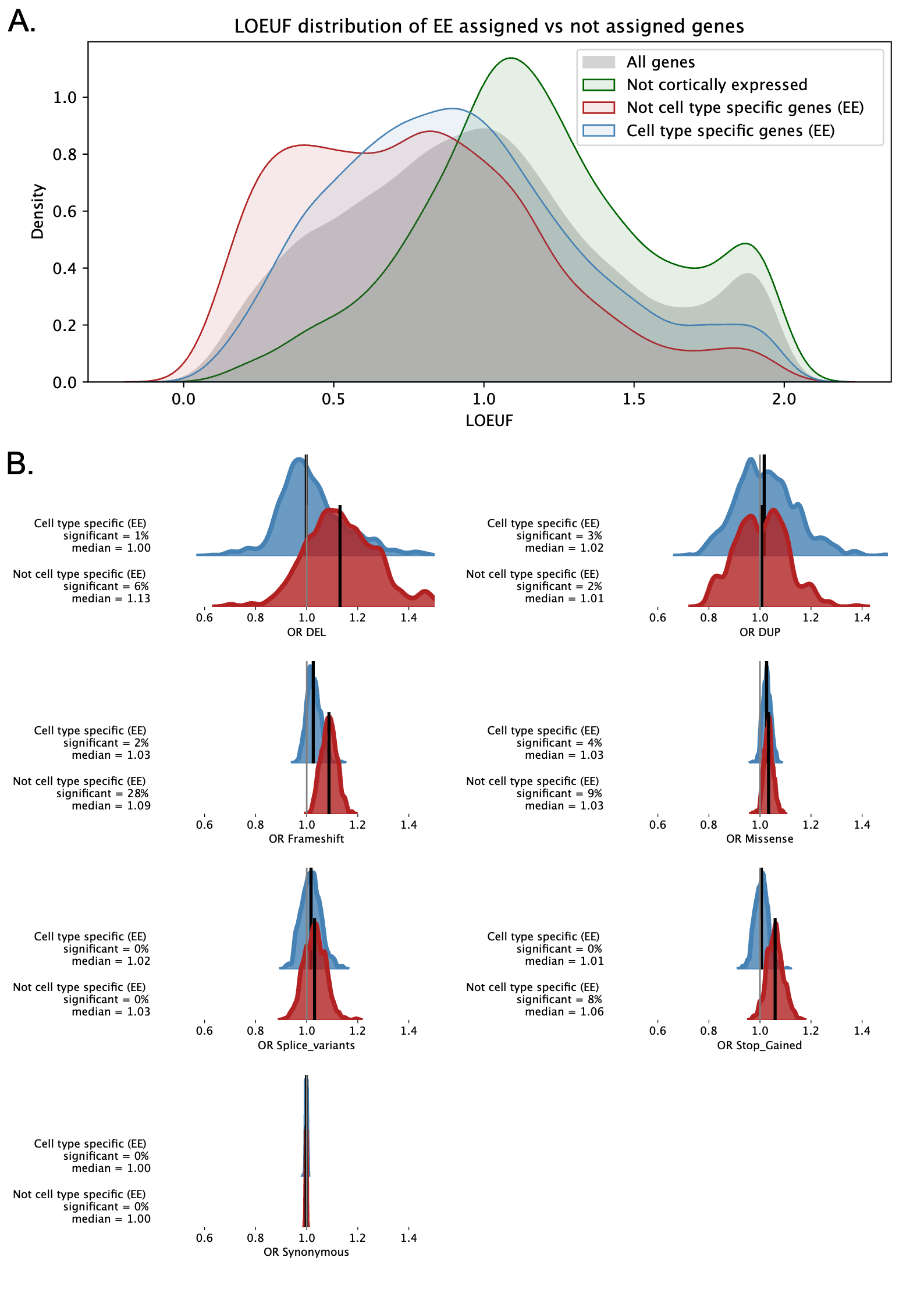

**Supplementary Figure 17. ASD association of the cell-type-specific genes assigned by Expression Enrichment.** (A) LOEUF distribution of the autosomal protein-coding genes, the genes not expressed in the PFC, the genes expressed in the cortex but not assigned to any cell type, and the genes assigned to at least one cell type. (B) For the 7 classes of variants, distributions of the ASD associations (odd ratios) were computed for 1,000 bootstraps of the genes expressed in the cortex but not assigned to any cell type (red) and the genes assigned to at least one cell type (blue).

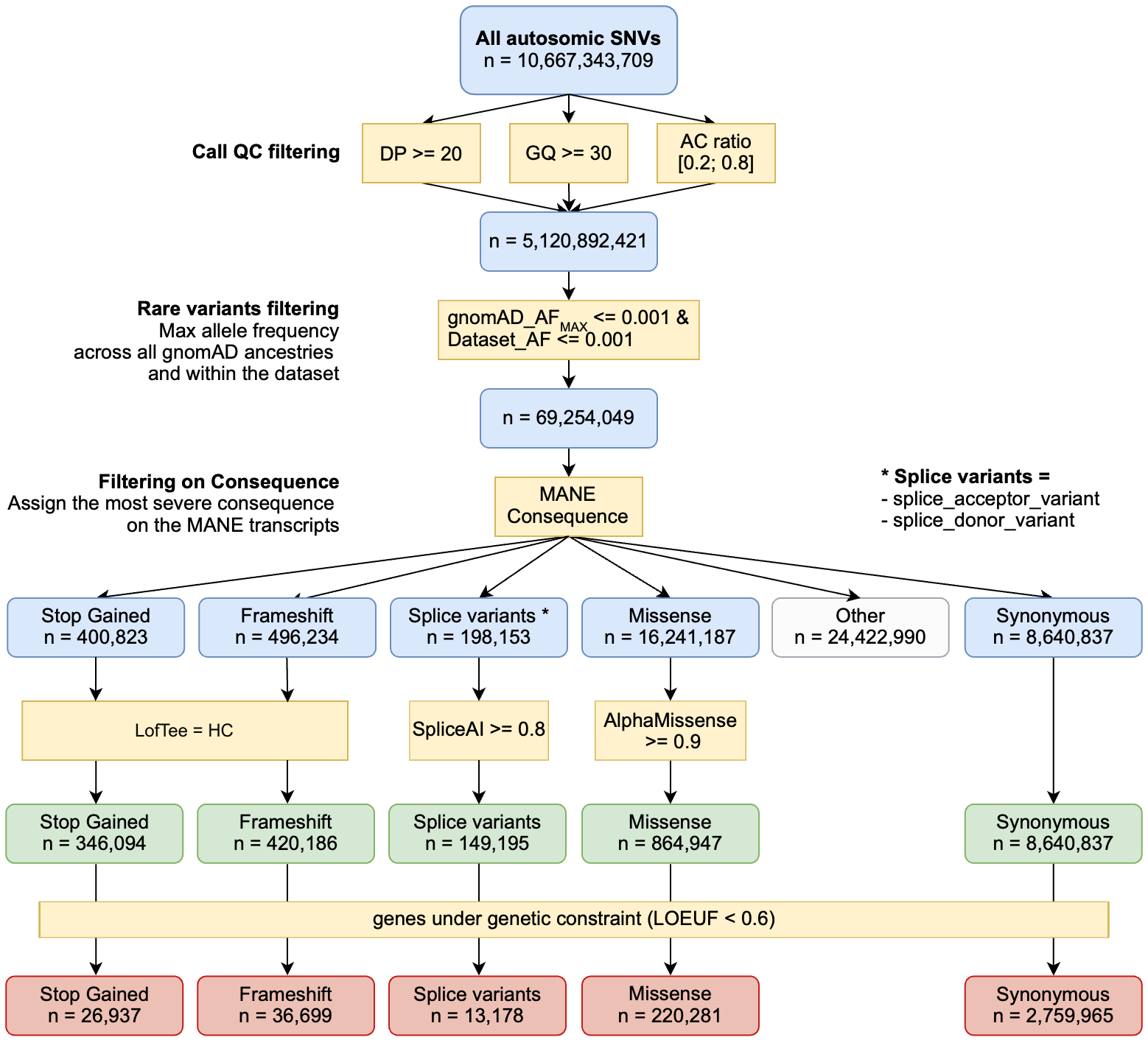

**Supplementary Figure 18. Total number of SNVs used in the analysis and the applied QC filtering.**

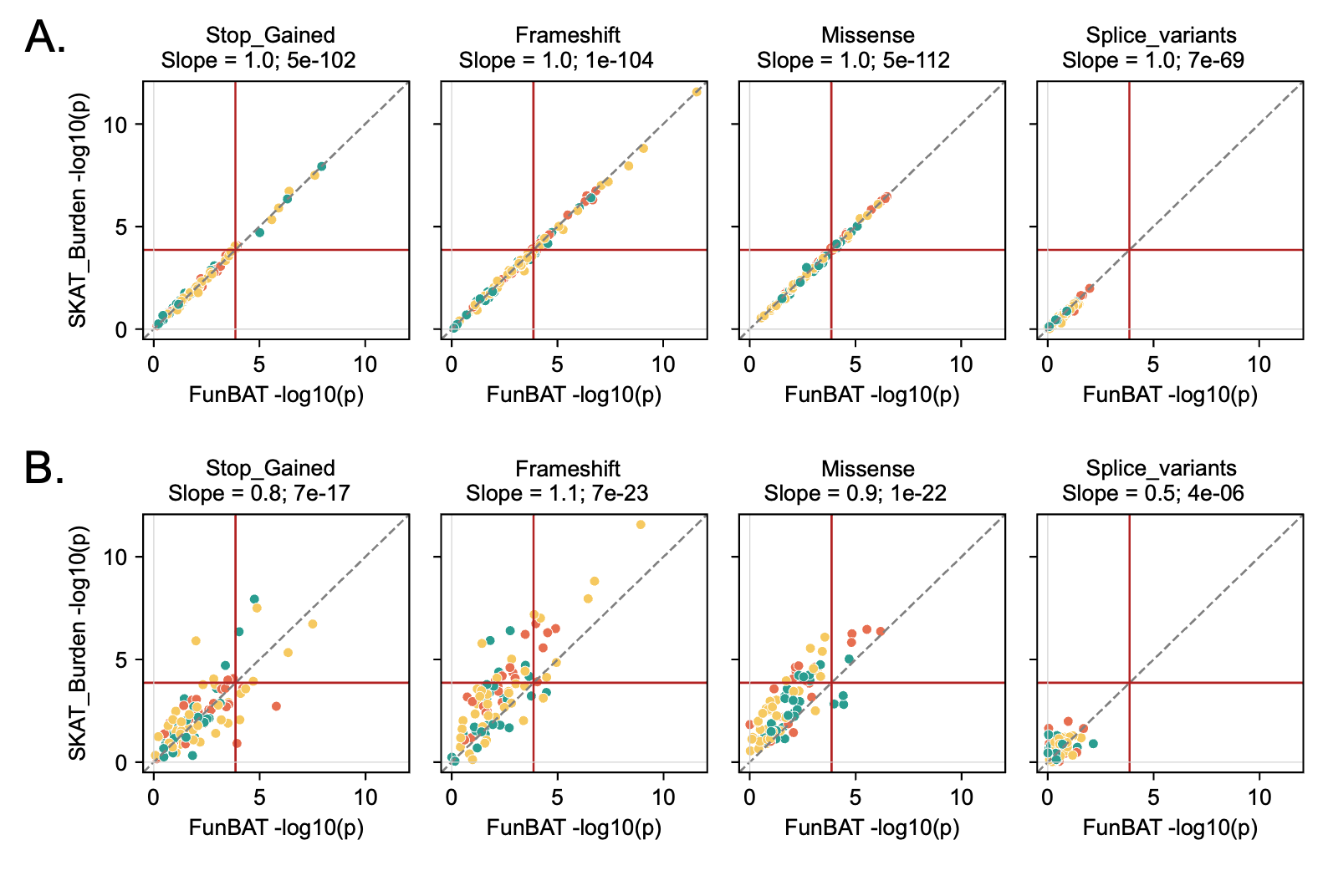

**Supplementary Figure 19. Comparison between p-values computed with SKAT-Burden and FunBAT.** Concordance analysis between the p-values (-log10) of the associations provided by SKAT-Burden and FunBAT. (A) FunBAT association of the cell types with the ASD risk computed through a logistic regression model and (B) FunBAT association computed with the conditional logistic regression model (model used for the main analyses) to correct for the dependence of the individuals. Data points correspond to the 91 cell types. The bias is reported for each class of variants as the slope of the regression line. The red lines represent the thresholds of significance (p=1e-4; Bonferroni correction, n = 364 tests).

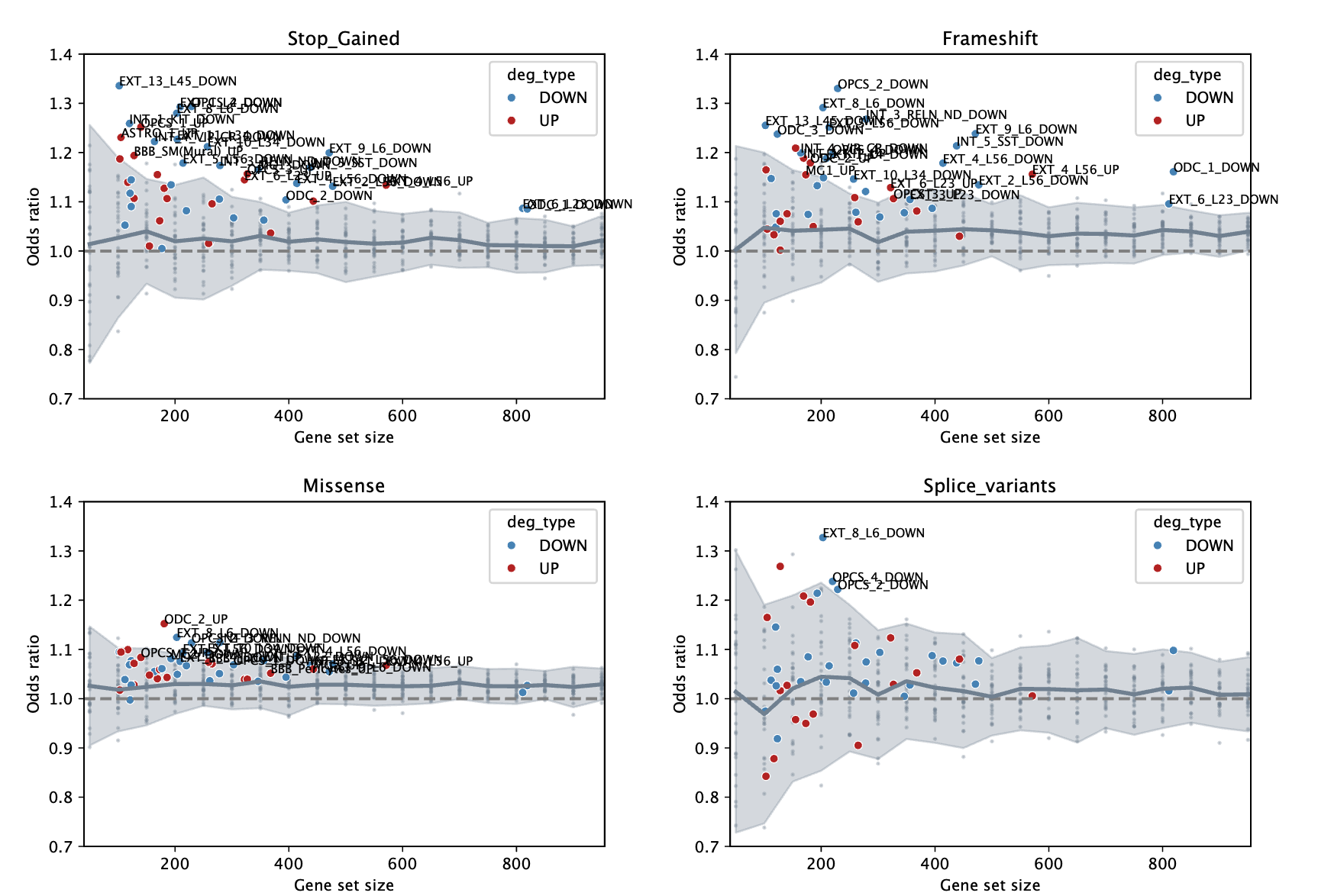

**Supplementary Figure 20. Distribution of ASD associations across random gene sets of multiple sizes.** Grey areas represent the IC95 of the ASD associations (odd ratios) computed on gene sets randomly created. Sizes of the random gene set vary from 50 genes to 900 genes to capture the whole size diversity of the ASD-DE gene sets. ASD associations of the ASD-DE gene sets were mapped to their corresponding gene set size and color coded with downregulated in blue and upregulated in red. ASD-DE gene sets with a significant ASD association (FDR < 0.05) were labeled.

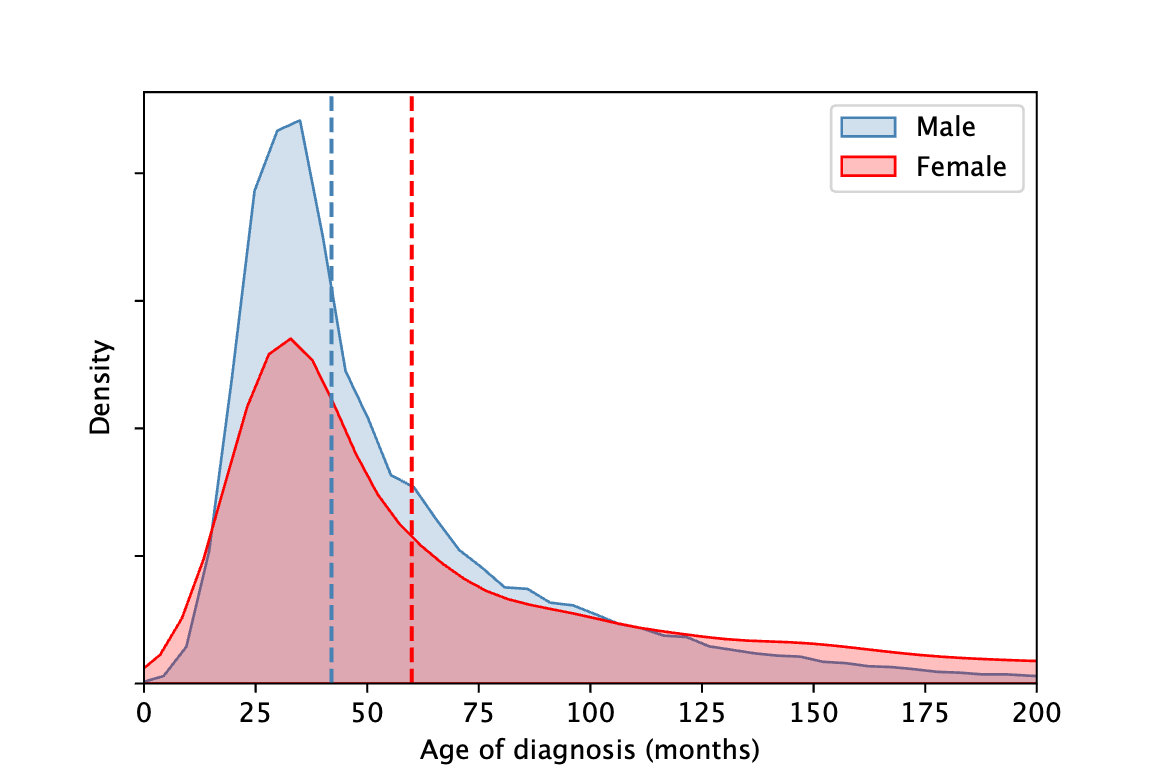

**Supplementary Figure 21. Distribution of age at diagnosis between males and females.** Medians for males and females are equal to 44 and 62 months, respectively. Mann-Whitney U-test p-value < 1e-300

### Supplementary tables

**Supplementary Table 1. Number of individuals with whole exome sequencing and used for the FunBAT analysis**

**Supplementary Table 2. Genetic enrichment of the 59 candidate genes identified in L56_TLE4 deep layer excitatory neurons.**

**Supplementary Table 3. Genetic enrichment of the 45 candidate genes identified in ID2 inhibitory neurons.**

**Supplementary Table 4. Genetic enrichment of the 55 candidate genes identified in microglia.**

**Supplementary Table 5. Equivalence table of the cell type annotations between the Herring et al. and the Wamsley et al. datasets.**
